# The mechanistic rules for species coexistence

**DOI:** 10.1101/2024.05.07.592948

**Authors:** Zhijie Zhang, Lutz Becks

**Affiliations:** Aquatic Ecology and Evolution, Limnological Institute, University of Konstanz, Konstanz, D-78464, Germany

## Abstract

Global biodiversity loss requires effective ecological community management, reliant on accurate predictions of species coexistence and community assembly^1-4^. Traditional methods, predicting coexistence from the effect of one species on another^5-8^ (i.e., phenomenologically), are sensitive to environmental context^9-13^. This is because they ignore the fundamental processes that can be applied across environments. While mechanistic approaches offer promise^14-17^, empirical tests remain rare^18,19^. Here, we integrated a mechanistic consumer-resource model with the growth of 12 phytoplankton species in monoculture over a range of phosphorous, nitrate or ammonium concentrations. We found that the mechanistic approach accurately predicts the composition of 960 communities across species richness and resource conditions. As confirmed by simulations, species competing for substitutable resources (ammonium vs. nitrate) exhibit greater diversity than those competing for essential resources (phosphorus vs. nitrate), especially when initial species richness is high. This is because when competing for essential resources, each species is likely to consume less of the resource that more limit its growth, violating the mechanistic rule of coexistence (each species must consume more of the resource that more limit it^16^). Our study highlights the power of the mechanistic approach in understanding and predicting species loss across environments and, ultimately, mitigating its pace.

---

Understanding and predicting species coexistence often relies on a phenomenological approach. This approach, inspired by the pioneering work of Lotka^5^ and Volterra^6^, infers competitive interactions from the negative effects of one species on the growth of another. Through this method, we have made huge strides in understanding coexistence. For example, it has revealed the relative strength of intra-to interspecific competition as a major determinant of species coexistence^20^ and recent studies have demonstrated its ability to predict community composition^7,8^. However, without an explicit statement regarding the causes of competition, it fails to fully capture the dependence of species coexistence on environments (e.g., resource availabilty^9-11^ and species richness^21^). This long-recognized limitation, dating back to Lotka and Volterra, has constrained the general applications of the phenomenological approach.

Consequently, mechanistic approaches are much needed. One such approach, formularized by MacArthur, describes how species consume and convert shared resources and thus compete with each other^14,15^. Expanding upon this groundwork, two rules for species coexistence were identified by Tilman^16,17^ (Fig. 1a). First, each species must be limited by different resources. Otherwise, a “superior” species with the lowest resource requirements will outcompete others. Second, each species must consume more of the resource that more limits itself. Despite the enduring interest in this mechanistic approach, direct empirical studies are rare^18,19^. The few studies that have been conducted mainly focus on resource requirement or the effect of resource availability on species coexistence, leaving the important role of resource consumption largely unexplored (but see refs.^22-24^). Furthermore, most studies focus on species pairs and a single type of resource (e.g., only essential^22^ or substitutable^23^ resources). Consequently, a comprehensive understanding of the proportion of species that meet Tilman’s two rules for coexistence remains elusive.

**Figure 1.**
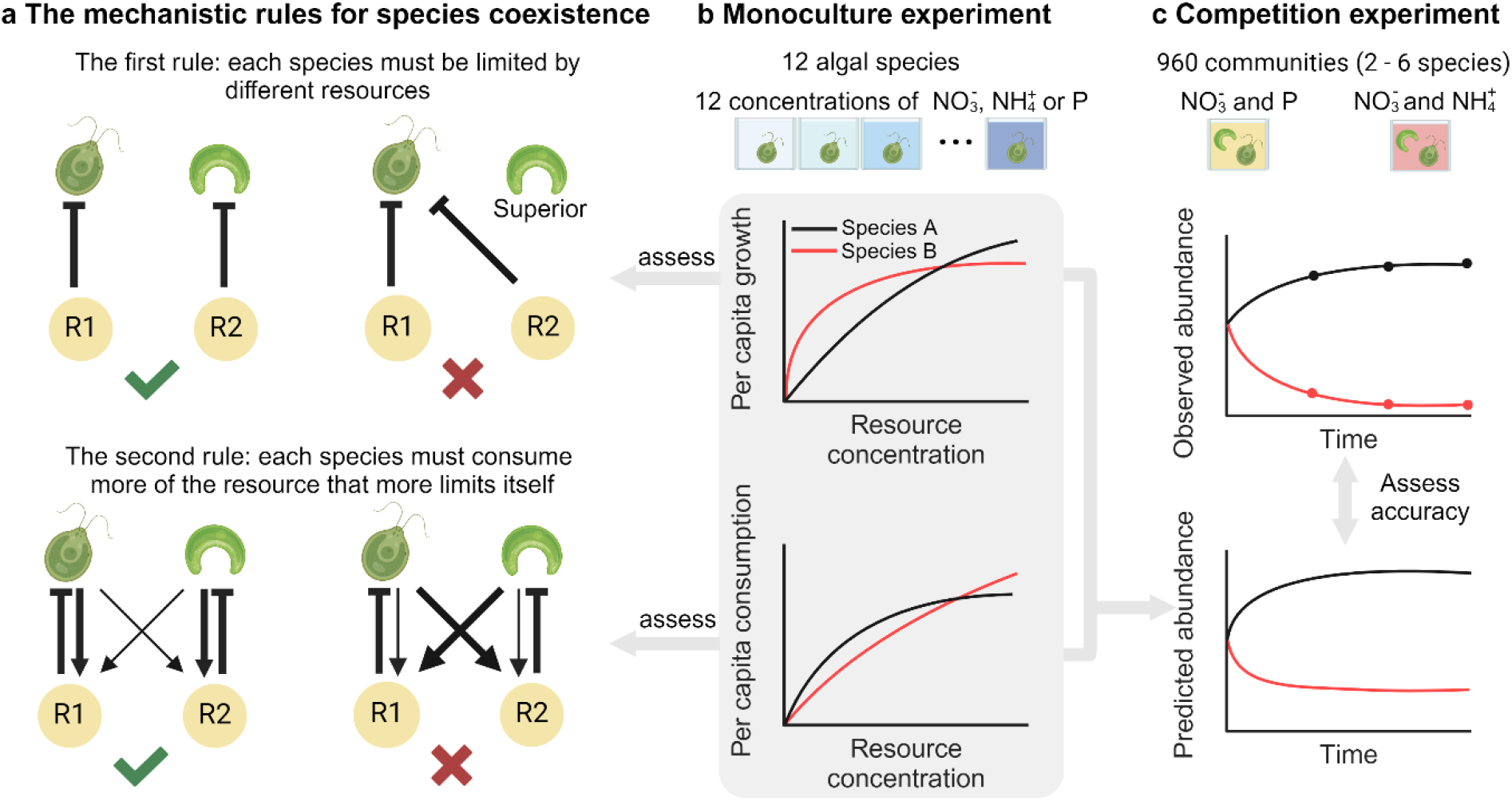
A mechanistic approach for understanding and predicting species coexistence. **a** Tilman’s rules for species coexistence, as illustrated by two algae consuming two resources. First, each species must be limited by different resources. Second, they must consume more of the resource that more limits itself. Inhibitors indicate the limitation of resources on algal growth (resource requirement). Arrows indicate the effect of algae on resources (resource consumption), with large width indicating large effect. **b** To quantify resource requirement and consumption, we combined a consumer-resource model with algal growth in monocultures under 12 concentrations of nitrate 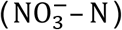, ammonium 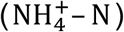, or phosphorus (P). We further assessed the proportion of species that met Tilman’s mechanistic rules. **c** We predicted community composition and assessed the predicative accuracy with competition experiments where two, three, four, or six species were competed for essential resources (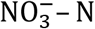 and P) or of two substitutable resources (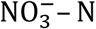 and 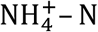).

Here, we test 1) whether the mechanistic approach can well-predict community composition and 2) to what extent the two rules for coexistence described by Tilman are fulfilled. We tracked the daily growth rates of 12 different phytoplankton species (green algae) in monocultures over four days (approx. zero to eight generations, depending on the resource concentrations). This was conducted under 12 concentrations of nitrate (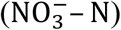), ammonium (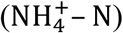), or phosphorus (P; Fig. 1b). To quantify the resource requirement and consumption for each species, we used Bayesian modelling to parameterize a consumer-resource model using the growth data and initial resource concentrations (see Methods for details). Next, we grew two, three, four, or six species in semi-continuous cultures where they competed for different ratios of two essential resources (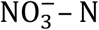 and P) or of two substitutable resources (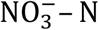 and 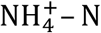). We tracked the community composition over 12 days with an automated pipeline that integrates high-content microscopy, imaging analysis, and machine learning; and compared the observed composition of these 960 communities with predictions from the consumer-resource model (Fig. 1b; Supplement S1: Fig. S1). Finally, we assessed the proportion of communities that meet Tilman’s two rules for coexistence with our experiment and simulation.

## The mechanistic approach predicts community composition

The per capita growth rate and consumption rate increased asymptotically with resource concentration for all 12 algal species when measured in monocultures, but we found large variation among species (Fig. 2a, b). Using this information on resource requirement and consumption, we asked whether the mechanistic, resource-consumer approach can predict the composition of 960 communities where species competed for essential or substitutable resources (Methods). As a control, we predicted the community composition (abundance) solely based on resource requirements. In this model, we assumed that the resource concentration was constant (i.e., no consumption from the algae). For essential resources (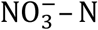 and P), we assumed that the growth of a species was determined by the resource that supports the lower growth rate, according to the Liebig’s law of minimum^25^. For substitutable resources (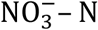 and 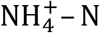), we assumed that the growth rate is the sum of the growth rates provided by both resources. Although the community composition strongly depended on resource condition (Supplement S2: Fig. S2), species abundance (as well as frequency; Supplement S3: Fig. S3) was well-predicted by resource requirement alone, with a mean accuracy of 67.8% (Fig. 2c). However, combining both resource requirement and consumption largely increased the predictive accuracy, reaching a mean accuracy of 86.4% (Fig. 2c; *F*_*1, 9484*_ = 6319.5, *P* < 0.001). This reveals that although resource requirement is a major determinant of species coexistence^26,27^, a precise prediction requires the bidirectional feedback between the species and its environment (e.g., resources).

**Figure 2.**
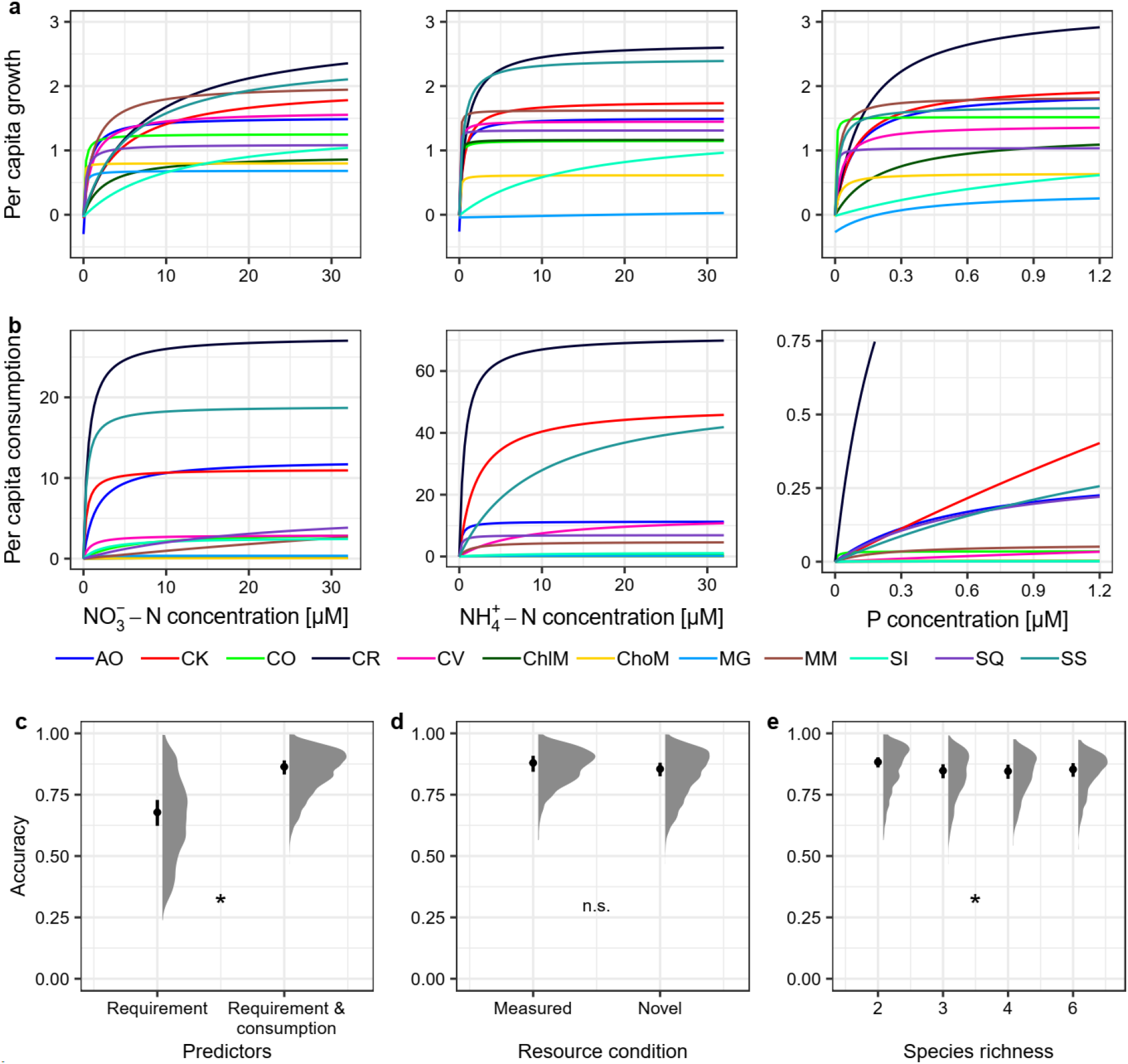
The resource-consumer model predicts community composition. **a, b** The resource requirement and consumption for nitrate 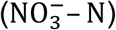, ammonium 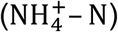, or phosphorus (P). Colors indicate different species (see Table S1 for the full names of the species). The curves were plotted using parameters fitted with the consumer-resource model using monoculture data (Methods). **c** We predicted the community composition with resource requirement alone (by assuming that resource concentration did not change over time) and with both resource requirement and consumption. **d** The predictive accuracy did not differ between the measured (same as the monoculture experiment) and novel conditions. **e** The predictive accuracy was slightly higher for two-species communities than multispecies communities. In **c, d, e**, error bars indicate the mean and 95% confidence intervals across competition experiments. Density plots indicate the distribution of the predictive accuracies of the 960 communities. Asterisks indicate significant differences between groups, as assessed by ANOVA F-test (*P* < 0.05).

Our mechanistic approach had robust predictive abilities across resource conditions (*F*_*1, 10*_ = 1.383, *P* = 0.267). It well-predicted community composition, not only in conditions where resource uses were measured but also in novel conditions (Fig. 2d). In comparison, previous work has found that the phenomenological approach, which infers competition from the negative effects of one species on the growth of another, was sensitive to abiotic environments, such as water and nutrient levels^9-11^. Such context-dependency makes it challenging to predict from one condition to another (but see ref.^12^ for a probabilistic approach). Furthermore, applying the phenomenological approach requires conducting experiments with 2^S^ -1 communities, where *S* is the species richness (but see refs.^7,28^ for approaches to reduce the experiment size). In contrast, the mechanistic approach sees the experiment size growing linearly with the species richness, as it requires measuring only monocultures. Since we found that the mechanistic approach also well-predicted community composition across species richness (Fig. 2e; mean accuracy > 84% for all species richness), it is now poised to be applied to biodiverse communities under various conditions.

Despite the generally accurate prediction, we noticed a slight decrease in accuracy with species richness (Fig. 2e; *F*_*3, 36*_ = 7.310, *P* < 0.001) and over time (Supplement S3; *F*_*4,4713*_ = 520.2, *P* < 0.001). Possibly, while we assumed that all the species consumed resources simultaneously, some may selectively consume one of the substitutable resources until its concentration becomes low. Furthermore, we found that alternative stable states^29^ were present, especially in diverse communities (Supplement S7). Therefore, the community may transition between these states with perturbation, which is possible since the cultures were refreshed every two days. Given the prevalence of disturbance and stochastic processes in natural communities, achieving accurate long-term predictions necessitates continuous monitoring.

## The mechanistic rules are more likely to be met when competing for substitutable resources

We next asked whether the species pairs can stably coexist by assessing Tilman’s two rules on species coexistence. We found that when competing for two essential resources (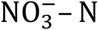 and P), only 30.3% of the species pairs met Tilman’s first rule for coexistence: each species must be limited by different resources (Fig. 3a). Among the species pairs that meet the first rule, only 40.0% of the pairs met the second: each species must consume more of the resource that more limits itself (Fig. 3b). Together, only 12.1% (8 out of 66) of the species pairs can stably coexist, which further depends on the resource supply^16^. When competing for substitutable resources (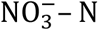 and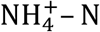), the probability of stable coexistence was higher (22.7%, 15 out of 66), with 37.9% and 60.0% of the pairs meeting the first and the second rules, respectively (Fig. 3). Both findings indicate the low probability of stable coexistence when species compete for two limiting resources, regardless of the resource type. Previous studies^30,31^ using phenomenological approaches revealed a comparable probability across taxonomic groups, ranging from 14.6% to 33.9%. However, because theory suggests multiple resources promote multispecies coexistence^32,33^, future study should explore its effect.

**Figure 3.**
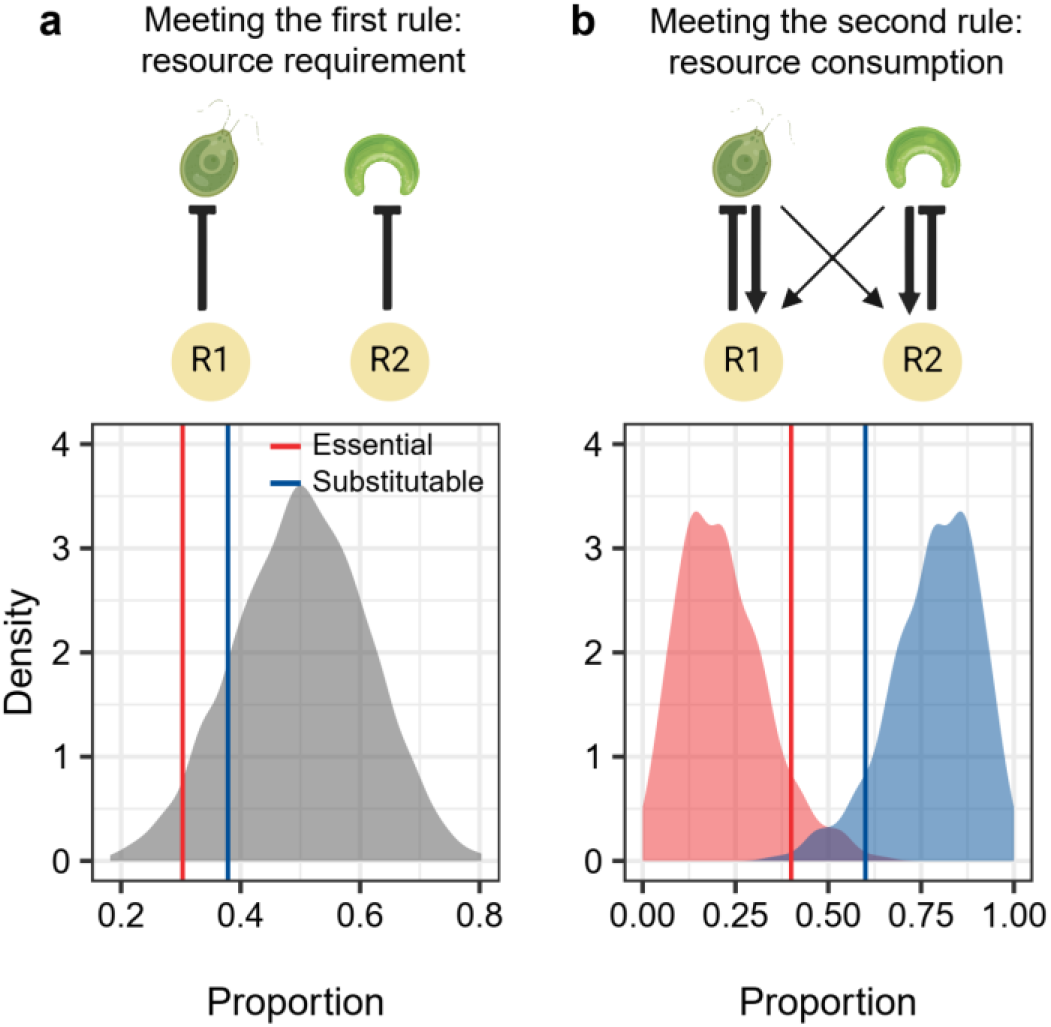
The mechanistic rules are more likely to be met when two species compete for substitutable resources. Vertical lines indicate the proportion calculated from the experiment. Density plots indicate the distribution of the proportion calculated from the simulation (i.e., the theoretical expectations). **a** The simulation showed that the median proportion of two species meeting the first rule was 50%, irrespective of the resource type (essential or substitutable). The proportion found in the experiment was 30.3% and 37.9% for essential and substitutable resources, respectively. **b** The proportion of two species meeting the second rule was low when competing for essential resources (40.0% in the experiment and 20.5% in the simulation) but was high for substitutable resources (60.0% in the experiment and 79.5% in the simulation).

To test whether our experimental results align with theoretical expectations, and whether there is a difference between competition for essential resources and competition for substitutable resources, we simulated 12 species over 999 times. For each species, the parameters in the consumer-resource model were randomly drawn from unique distributions (Methods and Supplement S4.1). On average, 50% of the species pairs met Tilman’s first rule (Fig. 3a; lower and upper 2.5% quantiles: 28.7% and 71.2%). Although the resource type did not affect the percentage of species that met the first rule, it did affect the second rule (Fig. 3b). Specifically, we found that species competing for essential resources were less likely to meet the second rule than those competing for substitutable resources (average percentage 20.5% *vs*. 79.5%). Both findings confirmed the results of our experiments.

But why species that compete for essential resources are less likely to meet Tilman’s second rule? Given the intrinsic link between consumption and growth rates (Supplement S4.2), the resource that a species consumes more is often the same resource that supports higher growth rates, consequently more limiting the species (Supplement S5). However, this is only true for substitutable resource. For essential resources, Liebig’s law of minimum reverses the results. Specifically, the only limiting resource is the one that supports the lower growth rates and thus is often the one that the species consumes less (Supplement S5). This contrasts with Tilman’s second rule: a species must consume more of the resource that more limits its growth rate. As a result, our finding, along with other theoretical work^34,35^, explains the inherent challenges of species coexistence when competing for essential resources, a concept known as the paradox of plankton^36^. However, substitutable resource for phytoplankton, such as different forms of nitrogen and different light spectrums^37^, is prevalent under natural conditions. Furthermore, phytoplankton serve as substitutable resources to their predators, offering an alternative pathway for the coexistence of phytoplankton^38^.

Last, we scaled up to multispecies communities. We found at the end of the experiment, Shannon diversity increased with initial species richness (Fig. 4a; *F*_*3, 36*_ = 7.64, *P* < 0.001), especially when the species competed for substitutable resources (Fig. 4a; *F*_*3, 575*_ = 22.88, *P* < 0.001). Our simulation confirmed it: although no more than two species can coexist when competing for two resources^39^, the probability of coexistence of two species increased with initial species richness (Fig. 4b; *F*_*3*_ = 407.5, *P* < 0.001), especially when competing for substitutable resources (Fig. 4b; *F*_*3*_ = 21.54, *P* < 0.001). This is because meeting the first rule is almost guaranteed in multispecies communities (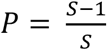, where *S* is the initial species richness; Supplement S6). Consequently, the probability of coexistence is predominately determined by the second rule, which is much more likely for substitutable resources.

**Figure 4.**
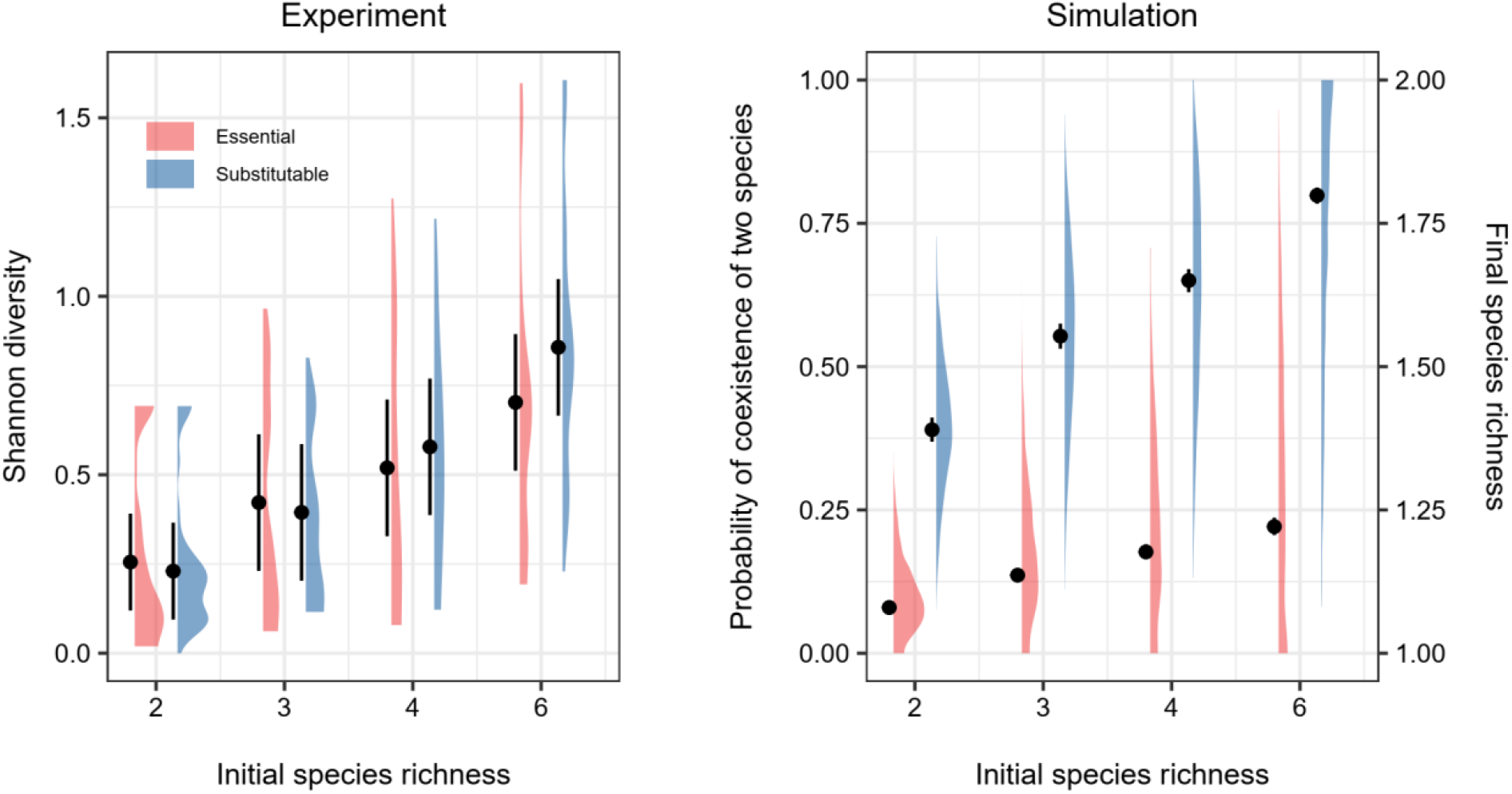
Diversity is promoted in multipspecies communities when species compete for substitutable resources. **a** At the end of the experiment, Shannon diversity increased with initial species richness, especially when the species competed for substitutable resources. **b** The simulation showed that the probability of coexistence of two species increased with initial species richness, especially when competing for substitutable resources. Density plots indicate the distribution of Shannon diversity of all communities in the experiment and that of probability of coexistence of two species (i.e., meeting both mechanistic rules) calculated from the simulation. Error bars indicate the means and 95% confidence intervals.

Our finding of the fundamental difference between essential and substitutable resources can advance our understanding of mechanisms governing species coexistence and evolution. For example, theoretical studies^40,41^ revealed that competition for substitutable resources led to character divergence^42^, while competition for essential resources led to character convergence. As suggested by our study, this is because stable coexistence through essential resources is unlikely and thus one way to achieve ‘neutral’ coexistence is by converging to ecologically identical species. Furthermore, our study suggests that the mechanistic model, which captures the processes underlying species interactions, can accurately predict community composition across a broad spectrum of abiotic and biotic conditions. This makes it a highly valuable tool for anticipating species loss and, ultimately, mitigating its pace.

## Methods

All the cultures were grown in TPP 96-well plates, at 20°C, constant light (65 µmol/m^2^/s), and static conditions.

### Study species

We selected 12 green algal species (Supplement S1: Table S1): *Acutodesmus obliquus* (abbreviation in Fig. 2: AO), *Chlamydomonas klinobasis* (CK), *Chlamydomonas oblonga (CO), Chlamydomonas reinhardtii* (CR), *Chlorella vulgaris* (CV), *Chlorella minutissima* (ChlM), *Choricystis minor* (ChoM), *Monoraphidium griffithii* (MG), *Monoraphidium minutum* (MM), *Scenedesmus intermedius* (SI), *Scenedesmus quadricauda* (SQ), *Scenedesmus sp* (SS), All species are from the algae collection at the Limnological Institute of the University of Konstanz. Before the experiments, stocks of each species were spread on agar plates and grown for seven days. Then, we picked a single colony for each species and grew it in modified WC medium (512 uM of nitrate and 25.6 uM of phosphorus) for nine days. The use of single colonies aims to minimize the effect of evolution on coexistence.

Two days before the experiments, we collected 2 ml culture for each species and washed the remaining nitrate and phosphorus by performing two rounds of centrifuging at 3,000 G for 5 minutes, discarding approx. 1.9 ml supernatant and replacing it with modified WC medium that did not contain nitrate and phosphorus. After that, we counted each culture and diluted it to approx. 500 cells/μl with a modified WC medium that did not contain nitrate and phosphorus. We let the culture rest for two days to reduce the remaining nutrients in the medium or potential intracellular storage of nutrients.

### Quantifying resource use from monoculture experiments

On the day of the experiment, for each species, we added 10 ul of diluted culture (∼ 5,000 cells) into 260 μl of modified WC medium. This medium was modified to contain varying concentrations of either nitrate resources 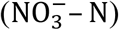, ammonium 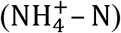, or phosphorus. For 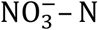, we reduced the nitrate concentration of the WC medium to one of the 12 levels: 0, 0.25, 0.5, 1, 2, 4, 8, 16, 32, 64, 128, and 512 μM. This ensured that nitrate was the major limiting factor, especially when low. For 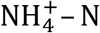, we replaced sodium nitrate with ammonium chloride and employed the same 12 concentrations. For P, we reduced the P concentration to one of the 12 levels: 0, 0.0125, 0.025, 0.05, 0.1, 0.2, 0.4, 0.8, 1.6, 3.2, 6.4 and 25.6 μM. With two replicates, we had a total of 864 cultures. All species and resource concentrations were randomized. We sampled each culture 20 μl after mixing for four consecutive days. Because our pilot experiment showed that most species experienced a lag phase of approx. 12 hours, we started our first sampling 12 hours after the inoculation. All culture preparation and sampling were done by a pipetting robot (OT-2, Opentrons). All samples were imaged and counted with a high-content microscopy (ImageXpress® Micro 4 High-Content Imaging System).

For a given species (*i*) and resource (*j*, 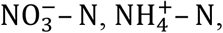, or P), we fit the growth data with a consumer-resource model that contains one species and one resource:

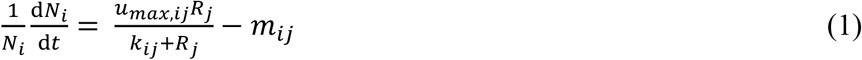

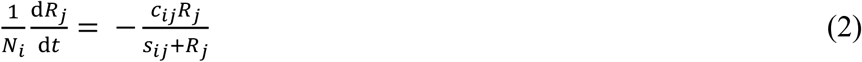

Equation (1) describes the per capita growth rate of species *i*, 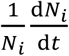, as a function of the concentration of resource *j* (*R*_*j*_), and thus indicates resource requirement. It follows the Monod equation^43^, where *u*_*max,ij*_ is the maximum birth rate of species *i* on resource *j, k*_*ij*_ is the half-saturation constant for birth rate (the concentration of resource *j* when species *i* reaches to half of its maximum birth rate), and *m*_*ij*_ is the mortality rate of species *i* in the absence of resource *j*. Equation (2) describes the per capita consumption rate of species *i* on resource *j*, 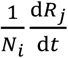, as a function of *R*_*j*_. It follows the type II functional response *sensu* Holling^44^, where *c*_*ij*_ is the maximum consumption rate, and *s*_*ij*_ is the half-saturation constant for consumption rate (the concentration of resource *j* when species *i* reaches to half of its maximum consumption rate).

While the theoretical model is continuous, abundance can only be measured at discrete times. Consequently, we calculated 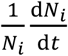 at time *t* as 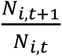, and 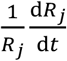 at time *t* as *ln* 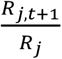, *w*here *t + 1* is one day after *t*. Theoretically, the relationship between resources and per capita growth rate (or consumption rate) can be linear. We tested this and found that the nonlinear relationship fitted the experimental data better (Supplement S8).

We applied a Bayesian approach to parameterize the model with the *brms* package^45^ in R language^46^. The resource consumption was quantified based on the combined information of per capita growth rate and initial resource concentrations (i.e., without the need for measuring resources over time). As a hypothetical example, consider a species growing in two media in which initial nitrate concentrations are 0 and 10 μM, respectively. While the initial per capita growth rate at the 10 μM concentration is high, it will decrease due to resource consumption and at a later time point reach the same rate as the initial per capita growth rate at the 0 μM concentration. Then, we can conclude that the amount of consumed nitrate is 10 μM. An additional assay in which we measured nitrate and phosphorus concentrations over time confirmed that our approach did well quantify the resource consumption (Supplement S9). To account for the non-independence of data from the same culture, we assumed that each culture had different maximum birth rates in the Bayesian model.

### Assessing community composition in competition experiments

In parallel with the monoculture experiment, we conducted competition experiments, where two, three, four, or six species were randomly chosen, mixed in the same ratio, and competed in modified WC media (20 μl culture added into 260 μl medium) for 12 days. We designed six resource conditions where the species competed for different ratios of essential resource (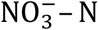 and P) and another six where the species competed for different ratios of substitutable resources (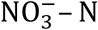 and 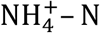 ; Fig. 1b). Note that we maintained high concentrations of other resources (e.g., potassium) while keeping the manipulated resources relatively low. This ensured that the manipulated resources were the major limiting resources. Four of the twelve resource conditions were used in the monoculture experiments (i.e., measured conditions), and the other eight were not (i.e., novel conditions; Fig. 1b). For species richness of two, we randomly selected 16 out of the 66 combinations of species. From these, we randomly selected eight combinations and then randomly added one, two, and four species, resulting in eight combinations each for species richness of three, four, and six, respectively. With two replicates, we had a total of 960 cultures. We sampled 20 μl of culture after mixing on day one, and then every two days starting from day four. The algal abundance for each sample was counted with the high-content microscopy (ImageXpress®), and species were identified with a machine-learning approach (Supplement S10). To maintain semi-continuous cultures, starting from day four, we mixed 160 ul of the old culture and 80 ul of fresh medium every two days.

### Predicting community composition with resource requirement and consumption

We predicted the community composition on day four to day twelve in the competition experiments from the resource requirement and consumption that were quantified from the monoculture experiments. The initial abundance of each species is based on the abundance on day one of the competition experiments. We modeled the abundance of each algal species and resource with a consumer-resource model that contains multispecies and two resources:

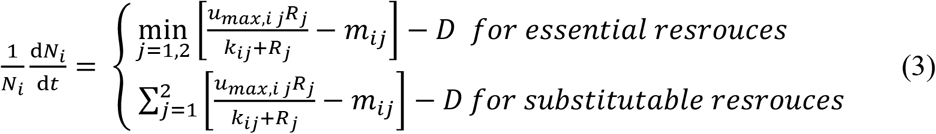

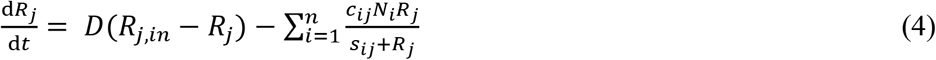

Equation (3) describes the per capita consumption rate of species *i* when grown on two resources. D is the dilution rate. It equals to ln 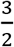 day^-1^ (160 ul old culture mixed with 80 ul fresh medium) at day 4, 6, 8, 10 and 12 and equals to 0 for the other days. All the other parameters follow the equation (1). For essential resources (here, 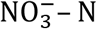 and P), the growth rate of the species is determined by the resource that supported a lower growth rate (i.e.,Liebig’s law). For substitutable resource (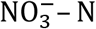 and 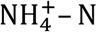), the growth rate is the sum of both growth rates. The choice of these two functions was further supported by an additional assay on growth rate under varying resource concentrations (Supplement S11). The equation (4) describes the change of resource *j*, as a function of *R*_*j*_ and *N*_*i*_. *D*(*R*_*j,in*_ *− R*_*j*_) describes the resources supply, with *R*_*j,in*_ representing input nutrient concentration (same as the initial nutrient concentration). 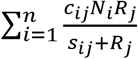 describes the resources consumption by all the species, with *n* representing the species richness in the community (i.e., *n* = 2, 3, 4, or 6). All the other parameters follow the equation (2). Note that, we transposed the equation (2) to simply equation (4). This resulted in the difference in the left-hand sides of equations (2) and (4) but did not affect the prediction.

In comparison, we also predicted the community composition with resource requirement alone, that is, assuming that the resource concentration does not change across the course of the experiment.

To assess whether our model well-predicts the community composition of competition experiments, for each community per time point, we calculated the predictive accuracy with a Bray–Curtis^47^ similarity index:

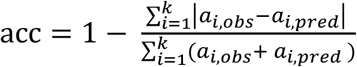

where *k, a*_*i,obs*_, *a*_*i,pred*_ are the number of species in the community, observed and predicted abundance of species *i*, respectively. The value of accuracy ranges from 0 to 1. A higher value indicates higher predictive accuracy, with a value of 1 indicating that the predicated abundance is exactly the same as the observed abundance. To give the rare species more weight (relative to their abundance) on the accuracy, we natural-log transformed the abundance after adding an abundance of one^48^. To minimize the effect of randomness on cell sampling and counting, we excluded data points where fewer than 10 cells were counted.

We first tested whether combining both resource requirement and consumption better predicted the community composition than using resource requirement alone. To do so, we conducted a linear mixed model with the *lme4* package^49^. The model included the predictive accuracy as the response variable and type of prediction (only resource requirement *vs*. both resource requirement and consumption) as the fixed effect. We then tested whether the predictive accuracy was consistent across time point, across novel and measured environments and across species richness. To do so, we conducted another linear mixed models. The model included the predictive accuracy (using both resource requirement and consumption) as the response variable and the time point (day four to day twelve), the type of medium (novel *vs*. measured) and species richness as the fixed effect. Both models included the identity of medium and species combination as the random effects. The predictive accuracy was logit-transformed to improve the normality of the residuals. The significance of the fixed effects was assessed with ANOVA.

### Assessing the proportion of species that meet Tilman’s rules for coexistence

#### Community of two consumers

To assess the proportion of species pairs that can stably coexist, we assessed Tilman’s two rules on species coexistence. To assess the proportion of species pairs that met Tilman’s first rule, each species must be limited by different resources, we tested whether the zero net growth isoclines (ZNGIs) of two species have a positive intersection (i.e., resource equilibrium) for each species pair (Supplement S4). This was done with the *sympy* library^50^ in python^51^.

For species pairs that met the first rule, we further assessed the proportion of species pairs that met Tilman’s second rule, each species must consume more of the resource that more limits itself. To do so, we determined for each species the resource that more limited itself. Mathematically, this is done by comparing the partial derivatives of the per capita growth rate of each species to each resource at the equilibrium, using the *math* library^52^. Then, we determined for each species the resource that it consumed more by comparing the per capita consumption rate of each species on each resource at the equilibrium.

To test whether our experimental results align with the theoretical expectation, we simulated 12 species. For each species, we randomly drew the parameters (e.g., *c*_*ij*_ and *s*_*ij*_) in the consumer-resource model from zero to one following a uniform distribution. Note that changing the range of the distribution (e.g., using from zero to two) will not affect the results (Supplement S5). Changing the type of the distribution (e.g., using Gaussian distribution) will affect the results quantitatively but not qualitatively (Supplement S5). We set the mortality rate to 0.1 for all the species. This is because our experiments showed that the species-specific mortality rate (mean: 0.01 day^-1^) is much lower than the mortality caused by dilution, which is constant.

Instead of modeling the per capita growth rate as a function of resource concentration, as we did above to quantify resource requirement, we modeled it as a function of per capita consumption rate. This is because although it is more practical to fit parameters with the former model; theoretically, an individual converts its consumed resource into growth rate^40^ (but see Supplement S4 for how the two methods match with each other). Specifically, we modeled the per capita growth rate of species *i* as:

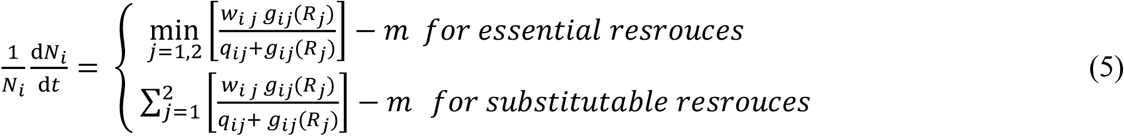

where *w*_*i,j*_ is a weighing factor, the value of one unit of the consumed resource *j* to species *i. g*_*ij*_ *(R*_*j*_) is per capita consumption rate of species *i* on resource *j*, i.e., 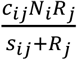, *q*_*ij*_ is the half-saturation constant for birth rate (the level of consumed resource *j* when species *i* reaches half of its maximum birth rate). m is the mortality rate. For the changes of resource, we modeled it as equation (4).

We only kept species that can grow when alone (i.e., had positive abundance when reaching to equilibrium), which meets the condition of our experiment. As above, we assessed the proportion of species pairs that met Tilman’s rules. We repeated the simulation over 999 times and calculated the average and 95% quantile for both proportions. Besides the simulation, we also derived the analytic solution in the linear system (Supplement S5), which confirmed the results of our simulation.

Note that as long as the two rules are met, the coexistence is further determined by the resource supply: the resource supply must fall within the region bounded by the consumption vectors of the two species (the third rule). This is explored in the supplement (Supplement S5.4).

#### Community of multiple consumers

While Tilman’s mechanistic rules are based on two consumers, we scaled it up to multiple consumers. The first rule should be adapted to: at least two of the species must be limited by different resources. In addition, the equilibrium of these two species must not be invadable by other species (i.e., all the other species have a negative growth rate at this equilibrium). Otherwise, at least one of the two species will be replaced by a third species. The second rule stays the same: for these two species, each species must consume more of the resource that more limits itself.

We calculated the mean probability of coexistence for each of the 999 sets of simulations above. For each simulation, we used all the combinations of two species (66), and randomly selected 99 combinations of three, four, or six species. Then, we tested whether the mean probability of coexistence of two species depended on the resource type and initial species richness with a linear model. The model included the mean probability of coexistence of each simulation as the response variable and the resource type, initial species richness, and their interaction as the explanatory variables.

In comparison, we tested with our competition experiment, whether the coexistence depended on the resource type and initial species richness with a linear-mixed effects model. The model included the Shannon diversity at the end of the experiment as the response variable; the resource type, initial species richness, and their interaction as the fixed effects; and the species combination (as we have two replicates) and resource condition as the random effects. Unlike the simulation, we used Shannon diversity. This is due to the duration of the competition experiment was not long enough to assess coexistence. However, by investigating Shannon diversity, we can still glean insights into whether the initial species richness and/or resource type promote coexistence.

## Acknowledgement

We thank Chuliang Song for help with deriving the analytic solution, Natascha Handke for helping the experiment, and Faruk Tök for conducting a pilot experiment. We also thank Toni Klauschies, Ville Mustonen, and Dietmar Straile for comments on the manuscript. ZZ was supported by DFG (ZH 113/2-1) and YSF co-funding at the University of Konstanz.

## Author contributions

Z.Z. and L.B. conceived the idea and designed the study. Z.Z. performed the experiment, analyzed the data, and conducted the simulation. Z.Z. drafted the manuscript and L.B. revised it.

## Data and materials availability

Should the manuscript be accepted, the data and code of the study will be archived in Figshare and the DOI will be included at the end of the article.

