## Supplementary material for "The mechanistic rules for species coexistence": SI

#### Supporting information

Zhijie Zhang, Lutz Becks

May 7, 2024

### Supplement S1 Experimental system

This section provides information on the species, and the species combination of the competition experiment.

**Table S1: The study species.** The abbreviation uses the first (or first three) letters of the genus and species.

| species name | Abbreviatoin | ID in the R code | Source* |
|---|---|---|---|
| *Acutodesmus obliquus* | AO | 13 | SAG 276-3a |
| *Chlamydomonas klinobasis* | CK | 56 | Konstanz |
| *Chlamydomonas oblonga* | CO | 7 | SAG 11-18a |
| *Chlamydomonas reinhardtii* | CR | c | Chlamydomonas Resource Center |
| *Chlorella minutissima* | ChlM | 5 | UTEX 2219 |
| *Chlorella vulgaris* | CV | 65 | Konstanz |
| *Choricystis minor* | ChoM | 9 | Konstanz |
| *Monoraphidium griffithii* | MG | 38 | Konstanz |
| *Monoraphidium minutum* | MM | 40 | SAG 243-1 |
| *Scenedesmus intermedius* | SI | 14 | Konstanz |
| *Scenedesmus quadricauda* | SQ | 16 | Konstanz |
| *Scenedesmus sp.* | SS | 3 | Konstanz |

*SAG: Sammlung von Algenkulturen der Universität Göttingen (with strain number)
Konstanz: collection at the University of Konstanz
UTEX: Culture Collection of Algae at UT-Austin (with strain number)

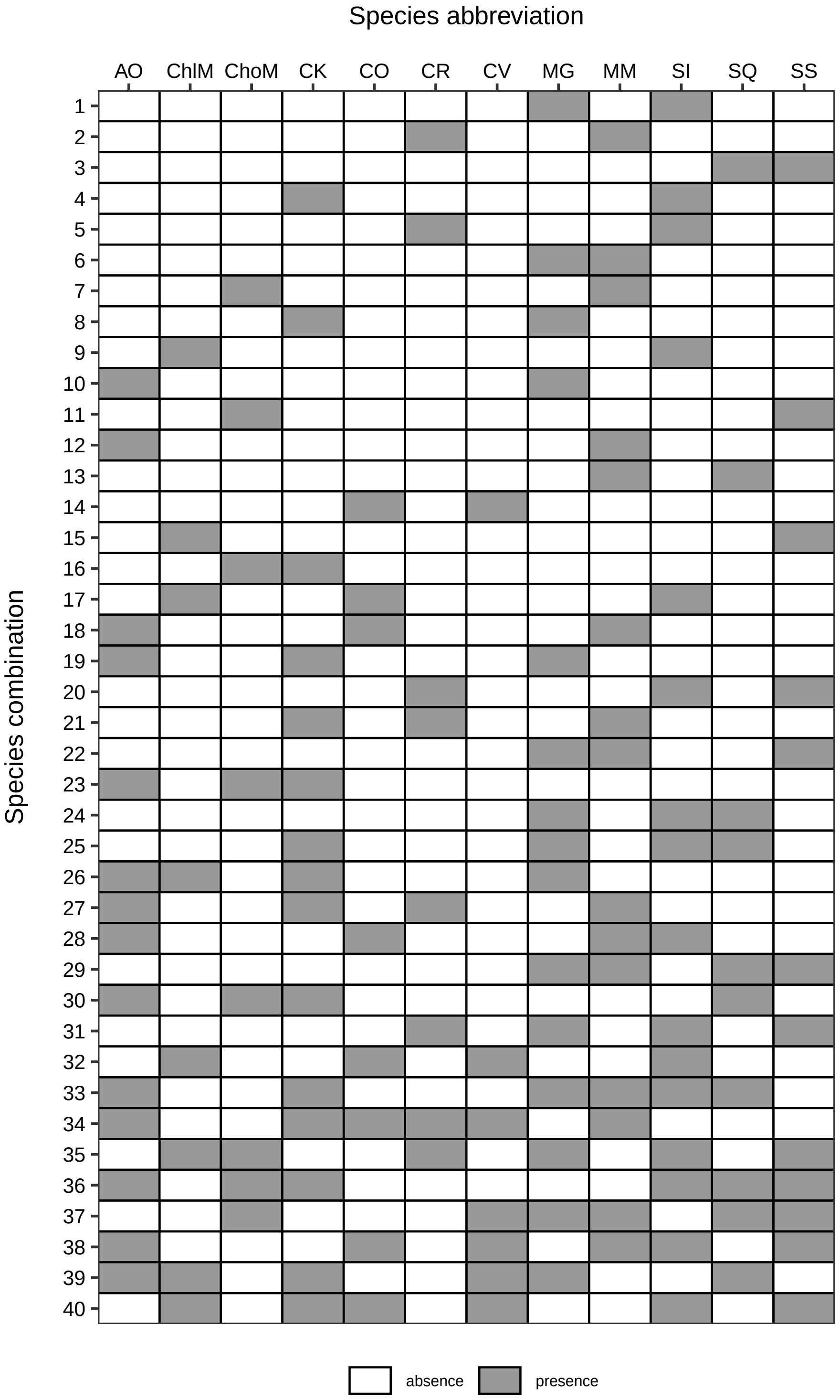


**Figure S1**: **The 40 species combinations of the competition experiments.** We randomly selected 16 combinations of species pairs. From these, we randomly selected eight combinations and then randomly added one, two, and four species, resulting in eight combinations each for species richness of three, four, and six, respectively. Grey cells indicate the presence of a certain species in the corresponding communities.

#### Supplement S2 The resource-dependency of the community composition

To test whether the community composition depends on resource conditions, we used permutational analysis of variance, as implemented in the *adonis2* function of the *vegan* package[1]. Specifically, for each species combination (Fig. S1), we used the species abundances at day 12 as the response variables and resource conditions as the explanatory variable. Same as the main text, we natural-log transformed the abundance after adding one.

We found that the community composition was significantly affect by resource condition for all of the 40 species combinations. Across the 40 combinations, the resource condition explained 87.5% of the variation among community composition (Fig. S2). This suggests that the community composition strongly depends on resource condition.

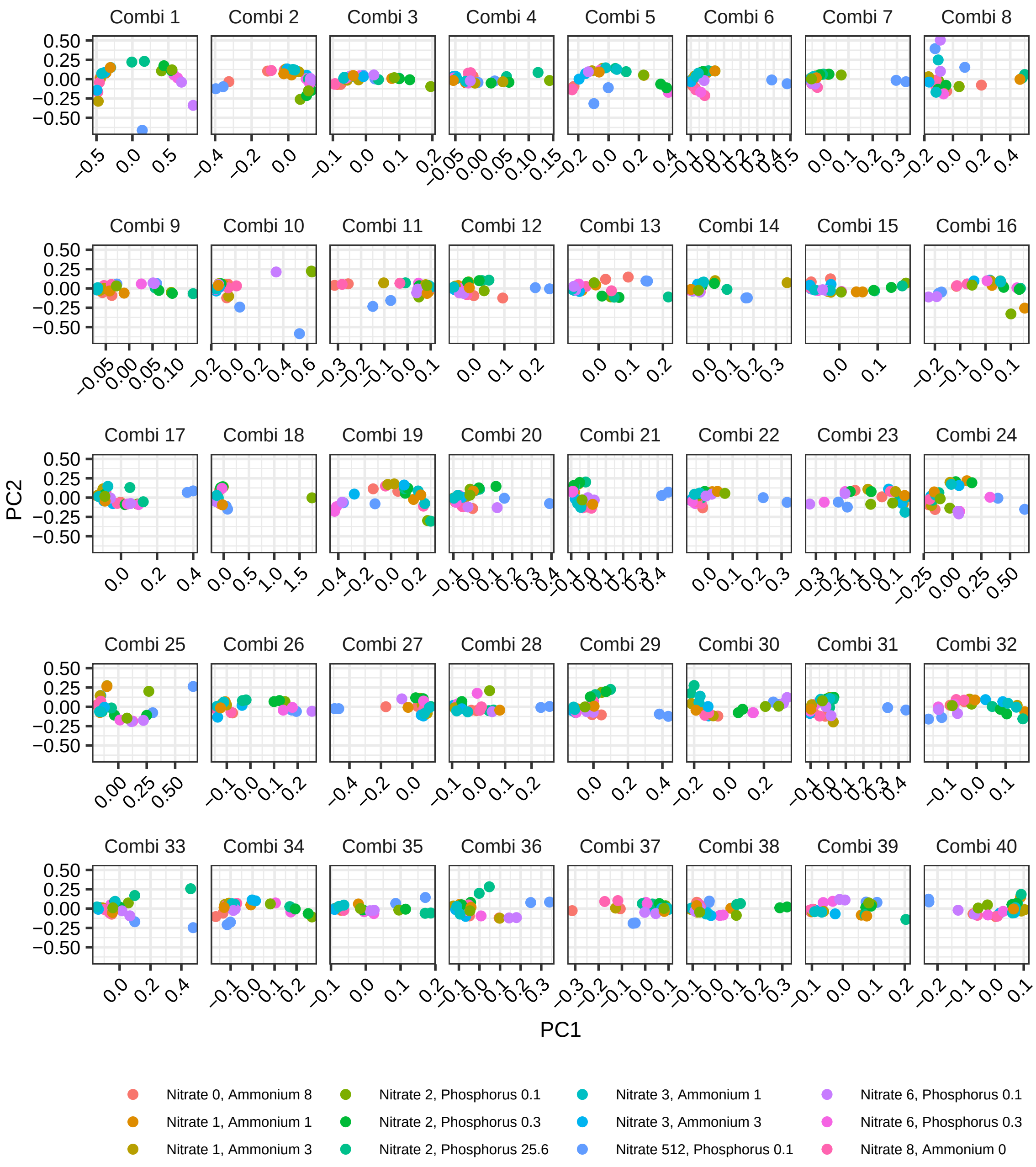


**Figure S2**: **The variation in the composition among communities.** For each species combination, we first computed the beta-similarity matrix for the all communities (12 resource conditions $\times$ 2 replicates). Then, we used Principal Component Analysis (PCA) to reduce the dimensionality of the beta matrix, and plotted the first two PCs. We can clearly see that the variation between replicates (same color) is low and that the variation between resource conditions (different colors) is large. The numbers after the resource indicate the resource concentration ($\mu M$). The details of the species combinations can be found in Fig. S1.

### Supplement S3  Additional results on predictive accuracy

While our main text focused on predicting species abundance, we also predicted the species frequency. Similarly, we assessed predictive accuracy on species frequency with a Bray–Curtis similarity index. Overall, the results showed the same pattern with the species abundance (Fig. S3). Combining both resource requirement and consumption resulted in higher predictive accuracy than using resource requirement alone ($F_{1,\ 9484} = 267.9, P < 0.001$). The accuracy did not significantly differ between measured and novel environments ($F_{1,\ 10} = 3.88, P = 0.077$), and between species richness ($F_{3,\ 36} = 2.14, P = 0.112$). While the accuracy differed between time points ($F_{1,\ 4713} = 11.67, P < 0.001$), it did not decrease with time. Instead, the accuracy at day 6 and day 8 was higher than that of day 4, probably because the between-species-variation in lag phase affected the early phase of the experiment.

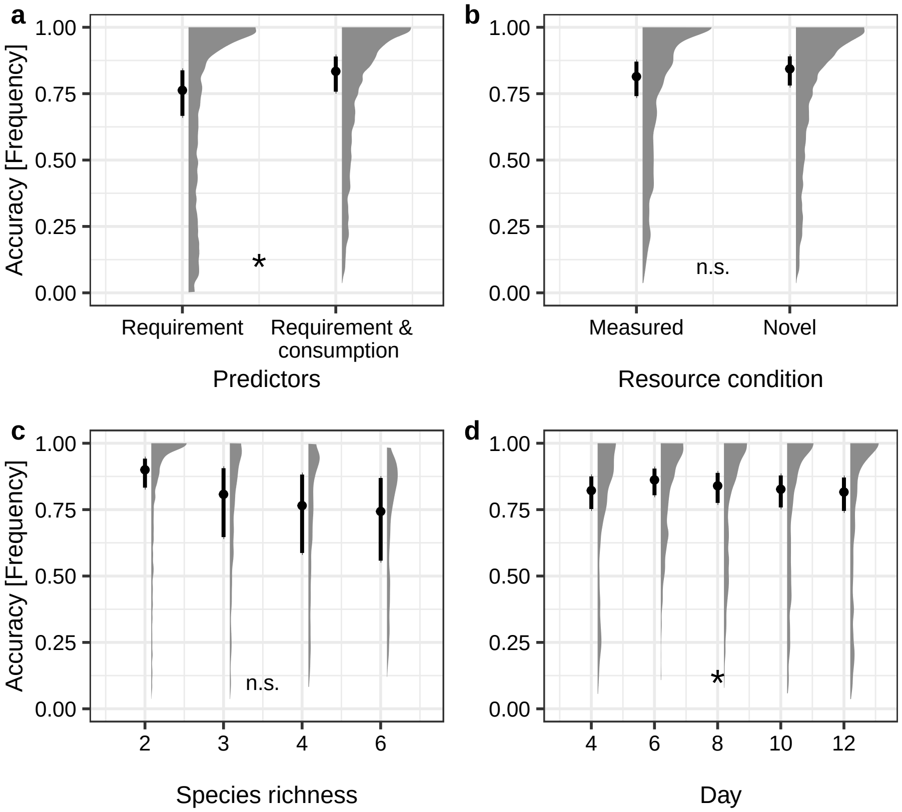


**Figure S3**: **The resource-consumer model predicts species frequency. a** We predicted the community composition with resource requirement alone and with both resource requirement and consumption. **b** The predictive accuracy did not differ between the measured (same as the monoculture experiment) and novel conditions. **c** The predictive accuracy did not differ between species richness. **d** The predictive accuracy was higher for days 6 and 8 than for day 4. Error bars indicate the means and 95% confidence intervals across competition experiments. Density plots indicate the distribution of the predictive accuracies of the 960 communities. Asterisks indicate significant differences between groups, as assessed by ANOVA F-test (P < 0.05).

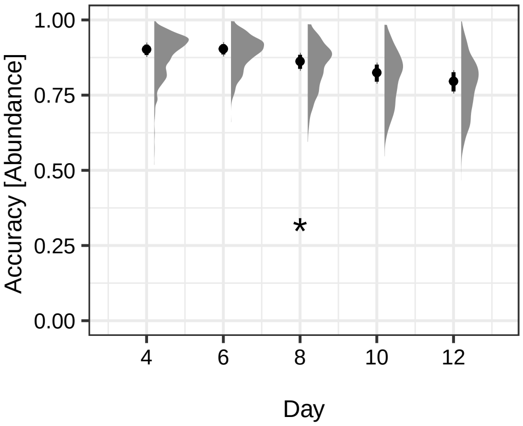


**Figure S4**: **The predictive accuracy on species abundance over time.** Density plots indicate the distribution of the predictive accuracies of the 960 communities. Asterisks indicate significant differences between groups, as assessed by ANOVA F-test ($P < 0.05$).

### Supplement S4 The model details

To enhance the clarity of this section, we did not specify *i* and *j*, that is, the identity of species and resources, respectively. Note that in latter sections, e.g., Supplement S5, we added *i* and *j* back as the subscripts.

#### Supplement S4.1 The model used in the simulation

##### Supplement S4.1.1 The general form

Biologically, an individual (e.g., an algal cell) first consumes a certain amount of resources according to the resource concentration:

$$\frac{1}{N}\frac{dR}{dt} = f(R) = -X \tag{S1}$$

where N and R are the abundance of the algae and the concentration of the resource, respectively. Then, it converses the consumed resource, *X*, to its growth rate according to:

$$\frac{1}{N}\frac{dN}{dt} = g(X) \tag{S2}$$

Substitute *X* with equation (S1), we get:

$$\begin{aligned}\frac{1}{N}\frac{dN}{dt} &= g(-f(R)) \\ &= h(R)\end{aligned} \tag{S3}$$

This reveals that the per capita growth rate, *g(X)*, can be translated to $h(R)$, a function of *R*.

##### Supplement S4.1.2 The linear form

While we used the nonlinear form in our simulation, here, we start with the linear form, which is more straightforward to understand. By assuming that the functions $f$ and $g$ are both linear, we have:

$$\frac{1}{N}\frac{dR}{dt} = f(R) = -cR = -X \tag{S4}$$

$$\frac{1}{N}\frac{dN}{dt} = g(X) = wX - m \tag{S5}$$

where $c$ is the consumption rate per resource unit, and $w$ is a weighing factor, the value of one unit of the consumed resource to the growth of the species. Substitute $X$ with equation (S4), we get:

$$\begin{aligned}\frac{1}{N}\frac{dN}{dt} &= g(X) \\ &= cwR - m \\ &= h(R)\end{aligned} \tag{S6}$$

##### Supplement S4.1.3 The nonlinear form (used in the simulation)

In the case of nonlinear consumption, same as equation (2) in the main text, we assume that an individual increases its resource consumption asymptotically with resource concentrations:

$$\frac{1}{N}\frac{dR}{dt} = -\frac{cR}{s+R} = -X \tag{S7}$$

Note that, here, the $c$ is the maximum consumption rate. Then, we assume that the resource conversion rate is nonlinear, that is, the growth rate increases asymptotically with consumed resource, $X$:

$$\begin{aligned}\frac{1}{N}\frac{dN}{dt} &= g(X) \\ &= \frac{wX}{q+X} - m\end{aligned} \tag{S8}$$

Substitute $X$ with equation (S7), we get:

$$\begin{aligned}\frac{1}{N}\frac{dN}{dt} &= \frac{w\frac{cR}{s+R}}{q+\frac{cR}{s+R}} - m \\ &= \frac{wcR}{qs+(c+q)R} - m \\ &= \frac{\frac{wc}{c+q}R}{\frac{qs}{c+q}+R} - m\end{aligned} \tag{S9}$$

By comparing equation (S9) and equation (1) in the main text, it is clear that

$$u_{max} = \frac{wc}{c+q}$$

and that

$$k = \frac{qs}{c+q}$$

This reveals that the theoretical (simulation) and empirical models are the same. However, it is easier to fit the experimental data with the empirical model, which is less complex despite the same number of parameters.

For each species, we randomly drew the parameters (e.g., $c_{ij}$ and $s_{ij}$) in the consumer-resource model from (0, 1) following a uniform distribution. We set the mortality rate to 0.1 for all the species. This is because our experiments showed that the species-specific mortality rate (mean: 0.01 day$^{-1}$) is much lower than the mortality caused by dilution, which is constant.

#### Supplement S4.2 The positive correlation between consumption and growth rates

Because the function $g$ is monotonic, it is clear that the per capita consumption rate and per capita growth rates are positively correlated (This is also revealed by equations S6 and S8). Biologically, this means that the more resources one species consumes, the faster it grows.

Additionally, we found that this was true in our experiment (Fig. S5)

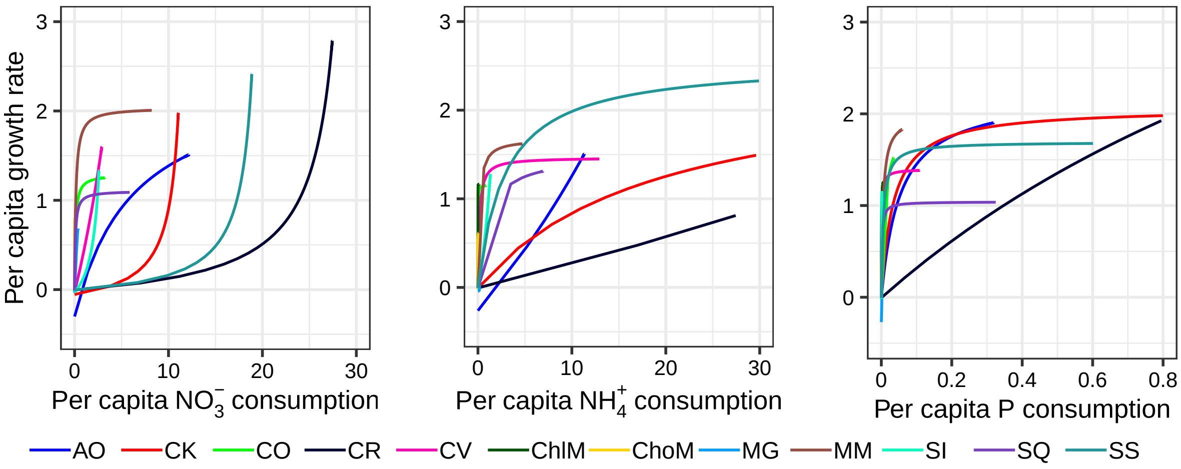


**Figure S5**: **The relationship between consumption and growth rates.** With the parameters fitted from the experiment, per capita growth rate and consumption rates were calculated across resource concentrations and plotted. Note that the species differed in their ranges of the x axis because they differed in their maximum consumption rates ($c_{ij}$)

### Supplement S5 The probability of two species meeting Tilman's rules

Here, we stick to the case where two species consume two resources. The general form is

$$\frac{dR_j}{dt} = a_j(R_j) - \sum_{i=1}^{2} N_i[f_{ij}(R_j)] \tag{S10}$$

$$\frac{dN_i}{dt} = N_i h_i(R_1, R_2) \tag{S11}$$

The two equations, (S10) and (S11), have the same form with equations (S1) and (S2). But there are multiple species and resources. In addition, we added subscripts to indicate the species (*i*) and resource (*j*). Last, we added supply rate, which is defined as a function of the resource concentration ($a_j(R_j)$), in equation (S10). Although, the effect of supply rate on species coexistence was not tested in our study, it can be incorporated for those who are interested in.

#### Supplement S5.1 The first rule: each species must be limited by different resources

As proved in Tilman 1980 [2], two species are limited by different resources when their zero net growth isoclines (ZNGIs) intersect. To draw the ZNGIs, let equation (S11) = 0.

##### Supplement S5.1.1 Linear system

To derive the analytic solution, let us start with the linear system. For the linear system, the first rule is equivalent to that each species has a lower minimum requirement(i.e., $R^*$) than the other in a certain resource, as illustrated by Fig. S6. We can also clearly see from the figure that the condition does not depend on the type of resources (essential *vs.* substitutable; Fig. S6).

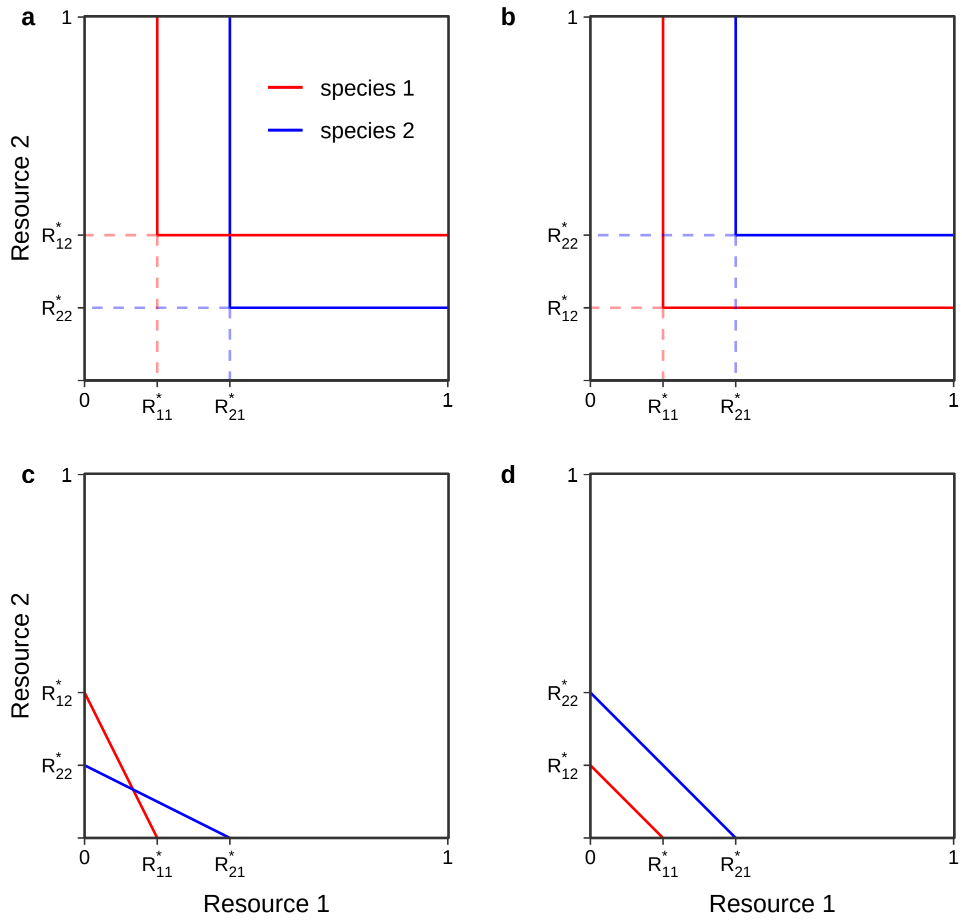


**Figure S6**: **Zero net growth isoclines (ZNGI) of two species competing for two resources (linear system). a & b**, two species are competing for two essential resources. **c & d**, two species are competing for two substitutable resources. In **a & c**, the ZNGIs (solid lines) intersect, indicating that the two species are limited by different resources (the first rule). Here, $R^*_{11} < R^*_{21}$ & $R^*_{12} > R^*_{22}$, where $R^*$ is the minimum resource requirement and the letters in the subscripts indicate the species (first letter) and resource (second letter). We used the same parameters for these two plots. In **b & d**, the ZNGIs do not intersect. Here, $R^*_{11} < R^*_{21}$ & $R^*_{12} < R^*_{22}$ . We used the same parameters for these two plots.

Let equation (S6) equal to 0, we can calculate the $R^*$ for each species per resource (Note that we used the same mortality for all species, as mentioned in the main text and at the end of Supplement S4.1.3):

$$R^*_{ij} = \frac{m}{c_{ij} w_{ij}} \tag{S12}$$

The two species will meet the first rule if

$$R^*_{11} < R^*_{21} \ \& \ R^*_{12} > R^*_{22} \text{ or } R^*_{11} > R^*_{21} \ \& \ R^*_{12} < R^*_{22}$$

Given S12 and the independence of the parameters (as they are drawn randomly), all $R_{ij}$ are independent with each other. Consequently,

$$P(R_{11}^* < R_{21}^* \mid R_{12}^* > R_{22}^*) = 0.5$$

that is, the probability of two species meeting the first rule is 0.5 in the linear system. In addition, we can see that the probability does not depend on the range or type of the distribution where we sample the parameters.

##### Supplement S5.1.2 Nonlinear system

For the model used in the main text (i.e., nonlinear system), we can calculate the $R^*$ for each species per resource according to equation (S9)

$$R_{ij}^* = \frac{m \cdot q_{ij} s_{ij}}{c_{ij} w_{ij} - (q_{ij} + c_{ij})m}$$

When competing for essential resources, the probability of two species meeting the first rule is still

$$P(R_{11}^* < R_{21}^* \mid R_{12}^* > R_{22}^*)$$

This is because the shape of the ZNGIs does not change with nonlinearity when competing for essential resources. Again, given that all the parameters are independent, the $R_{ij}$ are independent with each other. Consequently, the probability of two species meeting the first rule is still 0.5 in the nonlinear system.

When competing for substitutable resources, the condition changes slightly. This is because the nonlinearity of the ZNGIs adds complexity (Fig. S7).

First, when each species has a lower $R^*$ in one resource than the other species, the ZNGIs of the two species intersect (Fig. S7a). This is same as the linear system. Second, in most cases, when one species has lower $R^*$ in both resources than the other species, the ZNGIs do not intersect (Fig. S7b). Third, there are special cases where the ZNGIs intersect twice. In these cases, one species has higher $R^*$ in both resources than the other and its ZNGI is much more convex (Fig. S7c). However, because such cases are very rare (<0.5% according to simulation), we can conclude that the probability of two species meeting the first rule does not strongly depend on the type of resources, and approximately equals to 0.5.

For simplicity, we do not consider the cases where the ZNGIs intersect twice.

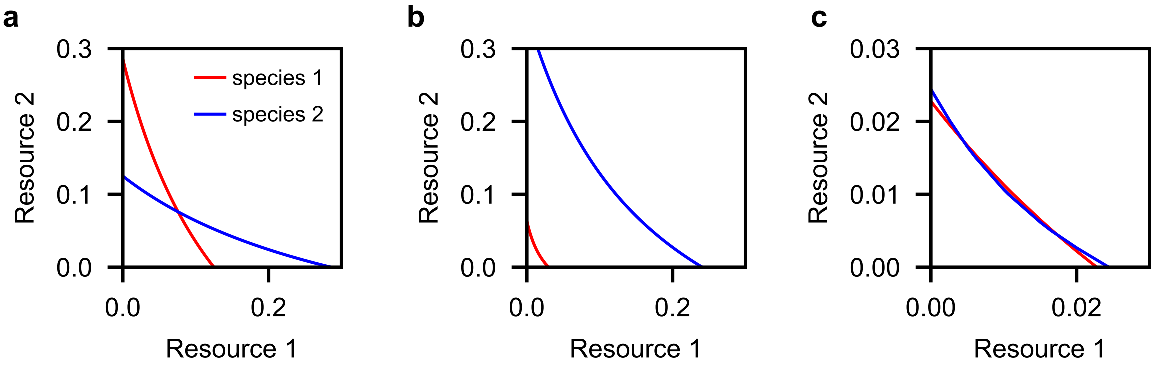


**Figure S7**: **Zero net growth isoclines (ZNGI) of two species competing for two substitutable resources (nonlinear system). a**, the ZNGIs (solid lines) intersect, indicating that the two species are limited by different resources (the first rule). Here, $R^*_{11} < R^*_{21}$ & $R^*_{12} > R^*_{22}$, where $R^*$ is the minimum resource requirement. **b**, the ZNGIs do not intersect. Here, $R^*_{11} < R^*_{21}$ & $R^*_{12} < R^*_{22}$. **c**, the ZNGIs intersect twice though $R^*_{11} < R^*_{21}$ & $R^*_{12} < R^*_{22}$.

#### Supplement S5.2 The second rule: each species must consume more of the resource that more limits itself

To assess whether the species pair meets the second rule, we need to assess at the equilibrium, 1) which species is more limited by which resource and 2) which species consumes which resource more.

For the general form, we assess these two questions with the two equations above: equations (S10) and (S11).

$$\frac{dR_j}{dt} = a_j(R_j) - \sum_{i=1}^{2} N_i[f_{ij}(R_j)] \tag{S10}$$

$$\frac{dN_i}{dt} = N_i h_i(R_1, R_2) \tag{S11}$$

To assess the first question, we calculate the following partial derivatives, all evaluated at equilibrium (*) with equation (S11)

$$\frac{\partial h_1}{\partial R_1^*};\quad \frac{\partial h_1}{\partial R_2^*};\quad \frac{\partial h_2}{\partial R_1^*};\quad \frac{\partial h_2}{\partial R_2^*}$$

Species 1 is more limited by resource 1 than species 2, if:

$$\frac{\partial h_1/\partial R_1^*}{\partial h_1/\partial R_2^*} > \frac{\partial h_2/\partial R_1^*}{\partial h_2/\partial R_2^*} \tag{I}$$

To assess the second question, we calculate all the per capita consumption rates, all evaluated at equilibrium (*) with equation (S10). Species 1 consumes more of resource 1 than species 2, if:

$$\frac{f_{11}(R_1^*)}{f_{12}(R_2^*)} > \frac{f_{21}(R_1^*)}{f_{22}(R_2^*)} \tag{II}$$

Consequently, the two species will meet the second rule if conditions (I) and (II) are both fulfilled or are both violated:

$$\frac{\partial h_1/\partial R_1^*}{\partial h_1/\partial R_2^*} > \frac{\partial h_2/\partial R_1^*}{\partial h_2/\partial R_2^*} \;\&\; \frac{f_{11}(R_1^*)}{f_{12}(R_2^*)} > \frac{f_{21}(R_1^*)}{f_{22}(R_2^*)}$$

or

$$\frac{\partial h_1/\partial R_1^*}{\partial h_1/\partial R_2^*} < \frac{\partial h_2/\partial R_1^*}{\partial h_2/\partial R_2^*} \;\&\; \frac{f_{11}(R_1^*)}{f_{12}(R_2^*)} < \frac{f_{21}(R_1^*)}{f_{22}(R_2^*)}$$

##### Supplement S5.2.1 Linear system

According to the section 1.1.2, we write the consumer-resource model:

The functions of resource consumption:

$$\frac{dR_1}{dt} = a_1(R_1) - c_{11}R_1N_1 - c_{21}R_1N_2 \tag{S13}$$
$$\frac{dR_2}{dt} = a_2(R_2) - c_{12}R_2N_1 - c_{22}R_2N_2 \tag{S14}$$

The functions of resource requirement when competing for essential resources :

$$\frac{dN_1}{dt} = N_1\ min[(c_{11}w_{11}R_1 - m),(c_{12}w_{12}R_2 - m)] \tag{S15}$$
$$\frac{dN_2}{dt} = N_2\ min[(c_{21}w_{21}R_1 - m),(c_{22}w_{22}R_2 - m)] \tag{S16}$$

The functions of resource requirement when competing for substitutable resources:

$$\frac{dN_1}{dt} = N_1(c_{11}w_{11}R_1 + c_{12}w_{12}R_2 - m) \tag{S17}$$
$$\frac{dN_2}{dt} = N_2(c_{21}w_{21}R_1 + c_{22}w_{22}R_2 - m) \tag{S18}$$

Because the prerequisite of the second rule is the first rule (i.e., the ZNGIs intersect), we assess 1) which species is limited by which resource and 2) which species consumes which resource more given that

$$R^*_{11} < R^*_{21} \ \&\ R^*_{12} > R^*_{22}$$

that is,

$$\frac{m}{c_{11}w_{11}} < \frac{m}{c_{21}w_{21}} \ \&\ \frac{m}{c_{12}w_{12}} > \frac{m}{c_{22}w_{22}}$$

which can be transposed to

$$\frac{c_{11}w_{11}}{c_{21}w_{21}} > 1 \ \&\ \frac{c_{12}w_{12}}{c_{22}w_{22}} < 1 \tag{S19}$$

**Assessing which species is more limited by which resource**

At first glance, this question seems to be equivalent to the first rule. However, fulfillment of the first rule does not immediately tell which species is more limited by which resource, and the latter further depends on the type of resources (essential *vs.* substitutable). We will explain this in the next paragraphs.

For <u>essential resources</u>, when evaluated at the equilibrium, the equations (S15) and (S16) can be simplified to:

$$\frac{dN_1}{dt} = N_1(c_{12}w_{12}R_2^* - m_1)$$
$$\frac{dN_2}{dt} = N_2(c_{21}w_{21}R_1^* - m_2)$$

We can see from the equations that the growth of species 1 is only affected by resource 2 at the equilibrium, the reverse is true for species 2. This indicates that species 1 is more limited by resource 2, as illustrated in Fig. S8a. A small change in resource 1 at the equilibrium will not affect the growth of species 1 (still on the ZNGI of species 1).

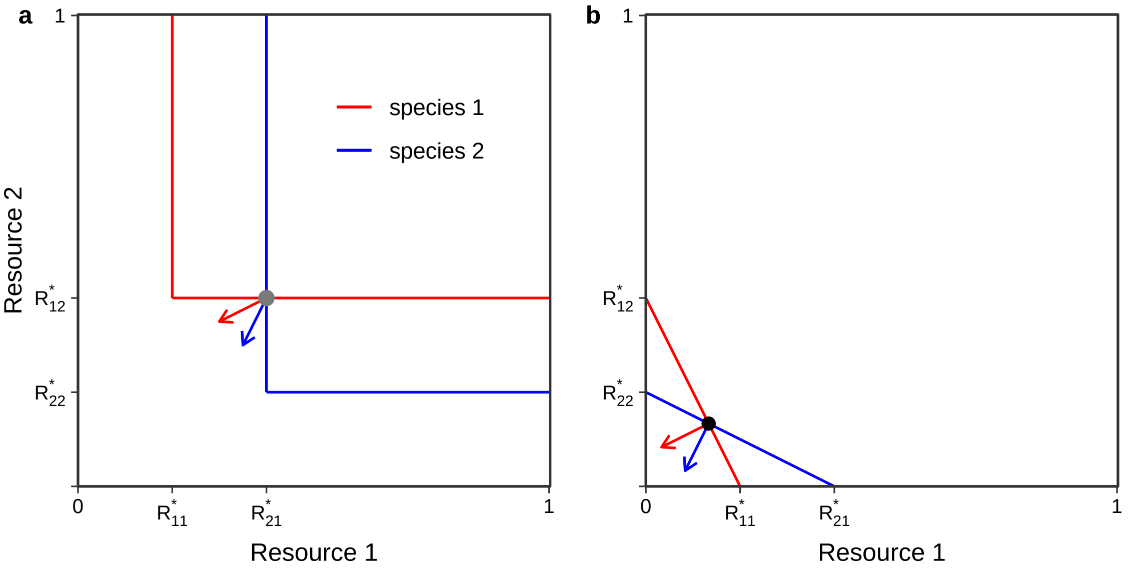


**Figure S8**: **Zero net growth isoclines (ZNGIs) and resource consumption of two species competing for two resources. a**, two species are competing for two essential resources. **b**, two species are competing for two substitutable resources. The arrows indicate the per capita consumption rate of the two species, with a flatter slope indicating higher consumption on resource 1. The two examples share the same parameters ($c_{11} = 0.5$, $c_{12} = 0.25$, $c_{21} = 0.5$, $c_{22} = 0.25$, $m_1 = m_2 = 0.1$, all other parameters equals to one). Consequently, $R^*_{11} < R^*_{21}$ & $R^*_{12} < R^*_{22}$. However, as explained in the text, when competing for essential resources, species 1 is more limited by resource 2. Consequently, the equilibrium (intersect) is unstable (gray), preventing coexistence. When competing for substitutable resources, species 1 is more limited by resource 1. Consequently, the equilibrium is stable (black)

Mathematically, we can check the following partial derivatives:

$$\frac{\partial h_1/\partial R_1^*}{\partial h_1/\partial R_2^*} = \frac{0}{c_{12}w_{12}}$$
$$\frac{\partial h_2/\partial R_1^*}{\partial h_2/\partial R_2^*} = \frac{c_{21}w_{21}}{0}$$

It is obvious that

$$\frac{\partial h_1/\partial R_1^*}{\partial h_1/\partial R_2^*} < \frac{\partial h_2/\partial R_1^*}{\partial h_2/\partial R_2^*} \tag{S20}$$

This confirms that species 1 is more limited by resource 2.

Similarly, for substitutable resources, we calculate the following partial derivatives according to equations (S17) and (S18):

$$\frac{\partial h_1/\partial R_1^*}{\partial h_1/\partial R_2^*} = \frac{c_{11}w_{11}}{c_{12}w_{12}}$$
$$\frac{\partial h_2/\partial R_1^*}{\partial h_2/\partial R_2^*} = \frac{c_{21}w_{21}}{c_{22}w_{22}}$$

Given the inequality (S19), it is obvious that

$$\frac{\partial h_1/\partial R_1^*}{\partial h_1/\partial R_2^*} > \frac{\partial h_2/\partial R_1^*}{\partial h_2/\partial R_2^*} \tag{S21}$$

This indicates that species 1 is more limited by resource 1, as illustrated in Fig. S8b. Note that this contrasts with the result when two species are competing for essential resources, that is, species 1 is more limited by resource 2 (compare S20 and S21). In other words, given the same parameters, as long as they meet the second rule for substitutable resources, they will not meet the second rule for essential resources..

**Assessing which species consumes which resource more**

We can calculate that

$$\frac{f_{11}(R_1^*)}{f_{12}(R_2^*)} = \frac{c_{11}R_1^*}{c_{12}R_2^*}$$
$$\frac{f_{21}(R_1^*)}{f_{22}(R_2^*)} = \frac{c_{21}R_1^*}{c_{22}R_2^*}$$

Consequently, the question of whether species 1 consumes resource 1 more,

$$\frac{f_{11}(R_1^*)}{f_{12}(R_2^*)} > \frac{f_{21}(R_1^*)}{f_{22}(R_2^*)} \tag{II}$$

is equivalent to whether

$$\frac{c_{11}}{c_{12}} > \frac{c_{21}}{c_{22}}$$

Given the inequality (S19),

$$\frac{c_{11}w_{11}}{c_{21}w_{21}} > 1 \,\&\, \frac{c_{12}w_{12}}{c_{22}w_{22}} < 1$$

It is obvious that

$$\frac{c_{11}w_{11}}{c_{21}w_{21}} > \frac{c_{12}w_{12}}{c_{22}w_{22}}$$

Let us consider first the special case, where

$$w_{11} = w_{21}$$

$$w_{12} = w_{22}$$

Then, we have

$$\frac{c_{11}}{c_{21}} > \frac{c_{12}}{c_{22}}$$

This guarantees condition (II), indicating that the probability of species 1 consuming resource 1 more is 1.00. Additionally, this probability does not depend on the type of resources. Taking the two questions together, our proof reveals that whether each species consumes more of the resource that more limits itself (the second rule) totally depends on the type of the resources. Specifically, when competing for essential resources, it is unlikely that the two species will meet the second rule. However, when competing for substitutable resources, it is very likely that they will meet the second rule (i.e., two species can stably coexist).

Solution for the general case

The next two pages are the proof of the probability. For those who are more interested in the result, the probability of two species meeting the second rule in linear system is $\frac{5}{6}$ when competing for substitutable resources and $\frac{1}{6}$ for essential resources.

For the probability of meeting the second rule when competing for substitutable resources, we need to calculate the probability of

$$\frac{c_{11}w_{11}}{c_{21}w_{21}} > \frac{c_{12}w_{12}}{c_{22}w_{22}}$$

given the inequality (S19),

$$\frac{c_{11}w_{11}}{c_{21}w_{21}} > 1 \ \& \ \frac{c_{12}w_{12}}{c_{22}w_{22}} < 1$$

To simplify the question, let us define a, b, c, and d, where

$$a = \frac{c_{11}}{c_{21}};\ b = \frac{c_{12}}{c_{22}};\ c = \frac{w_{21}}{w_{11}};\ d = \frac{w_{22}}{w_{11}}$$

Then, our question is equivalent to

$$P(a > b \mid a > c \ \& \ b < d)$$

So, we need to calculate the PDF of a, given that $a > c$; and the PDF of b, given that $b < d$. Because all parameters follow a uniform distribution ranging from 0 to 1, without any condition, the PDF of a, b, c, and d are:

$$f(x) = \begin{cases} \frac{1}{2} & \text{for } 0 < x < 1 \\ \frac{1}{2x^2} & \text{for } x \geq 1 \\ 0 & \text{otherwise} \end{cases}$$

See ref. [3] for details of calculating the PDF.

Then, we calculate the conditional CDF of $x_1 \leq a \leq x_1 + \delta$ given that $a > c$. According to Bayesian theorem,

$$\begin{aligned} P(x_1 \leq a \leq x_1 + \delta \mid a > c) &= \frac{P(a > c \mid x_1 \leq a \leq x_1 + \delta) \cdot P(x_1 \leq a \leq x_1 + \delta)}{P(a > c)} \\ &= \frac{\int_0^{x_1} f(x)\,dx \ \cdot f(x_1) \cdot \delta}{0.5} \end{aligned}$$

After calculating the integration, we get the PDF of $a$, given that $a > c$ :

$$f_{X_1}(x_1) = \frac{P(x_1 \leq a \leq x_1 + \delta \mid a > c)}{\delta} = \begin{cases} \frac{x_1}{2} & \text{for } 0 < x_1 < 1 \\ \frac{2x_1 - 1}{2x_1^3} & \text{for } x_1 \geq 1 \\ 0 & \text{otherwise} \end{cases}$$

Similarly, we do it for $b$.

For the CDF:

$$\begin{aligned} P(x_2 \leq b \leq x_2 + \delta \mid b < d) &= \frac{P(b < d \mid x_2 \leq b \leq x_2 + \delta) \cdot P(x_2 \leq b \leq x_2 + \delta)}{P(b < d)} \\ &= \frac{\int_{x_2}^{+\infty} f(x)\, dx \; \cdot f(x_2) \cdot \delta}{0.5} \end{aligned}$$

For the PDF:

$$f_{X_2}(x_2) = \begin{cases} \frac{2 - x_2}{2} & \text{for } 0 < x_2 < 1 \\ \frac{1}{2x_2^3} & \text{for } x_2 \geq 1 \\ 0 & \text{otherwise} \end{cases}$$

Finally,

$$\begin{aligned} P(a > b \mid a > c \,\&\, b < d) &= \int_0^{+\infty} F_{X_2}(x_1) f_{x_1}(x_1) dx_1 \\ &= \frac{5}{6} \end{aligned}$$

This is the probability of two species meeting the second rule (each species consumes more of the resources that more limits itself) when competing for substitutable resources. For essential resources, it is $P(a < b \mid a > c \,\&\, b < d)$, i.e., $\frac{1}{6}$.

##### Supplement S5.2.2 Nonlinear system

Similar to the linear system, we can write the consumer-resource models according to section 1.1.3. Again, we need to assess at the equilibrium, 1) which species is more limited by which resource and 2) which species consumes which resource more. We were not able to solve it analytically. However, the simulation showed that probability of two species meeting the second rule when competing for substitutable resources was 0.8, and 0.2 for essential resources. This is close to that of the linear system.

#### Supplement S5.3 Discussion

Our proof showed that provided the fulfillment of the first rule, the fulfillment of the second rule depends on the type of resources. We can see from above, that when competing for substitutable resources in the linear system, $\partial h_i / \partial R_j^*$, which describes the effect of resource *j* on species *i*, is proportional to $c_{ij}$, the consumption rate per resource unit. Consequently, each species is more likely to consume the resource that more limits itself. In contrast, when competing for essential resources, Liebig's law reverses the results. Specifically, the only limiting resource is the one that supports lower growth rates (i.e., a lower $c_{ij} w_{ij} R_j^*$), and thus is often the one that the species consume less (i.e., a lower $c_{ij} R_j^*$). Note that we assumed that the different parameters were independent in our models, violation of this assumption can increase the probability can increase or decrease the probability of stable coexistence [4].

The contrasting difference between essential and substitutable resources has even confused experts in this field (e.g., Fig. 2.8 in Chase & Leibold [5]; also see discussion in the last paragraph of section "Coexistence And Contemporary niche theory" in Letten et al [6]). Revealing this difference can deepen our understanding of mechanism underlying species coexistence.

#### Supplement S5.4 The third rule: the resource supply must fall within the region bounded by the consumption vectors of the two species

As proved in Tilman 1980 [2], as long as the first and second rules are met, whether two species coexist is determined by the resource supply vector, $\overrightarrow{U}$:

$$\overrightarrow{U} = [a_1(R_1), a_2(R_2)]$$

where $a_1(R_1)$ and $a_2(R_2)$ are the supplies of resources 1 and 2 in equation S10.

When the supply vector falls outside the region bounded by the consumption vectors (dashed lines in Fig. S9a, b), one species outcompetes the other. When the supply vector falls within the bounded region and the equilibrium is stable (second rule met), two species coexist (Fig. S9a). When the equilibrium is not stable, which species outcompetes the other depends on the initial density (i.e., priority effect; Fig. S9b)

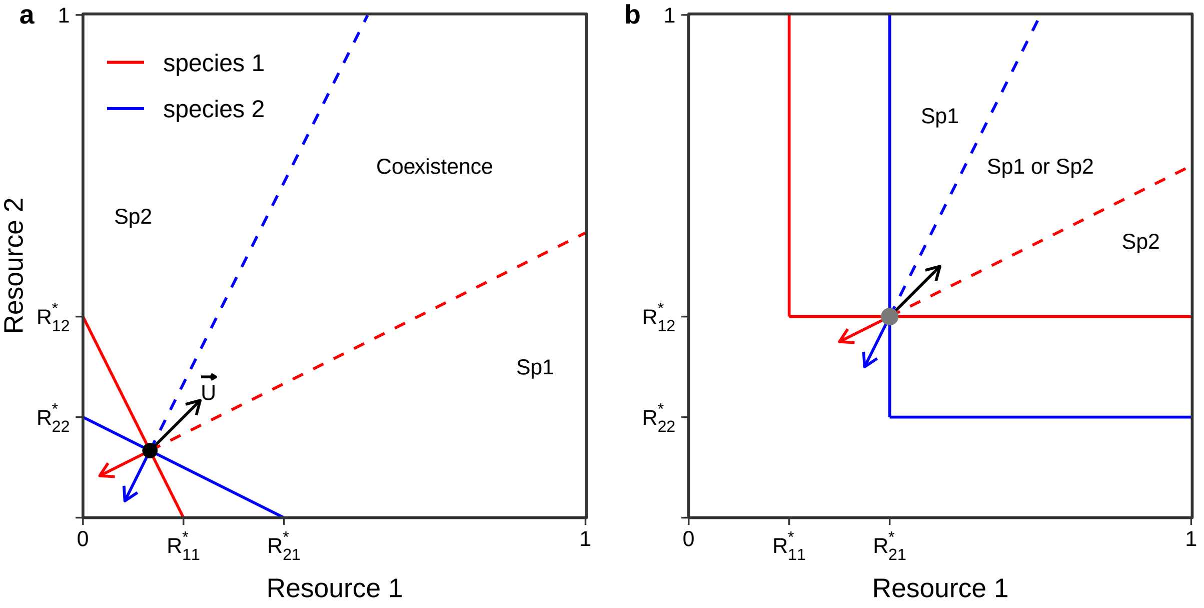


**Figure S9**: **Resource supply vector determines the outcomes of the two species.** **a** Two species coexist when the equilibrium is stable (both first and second rules met; black dot) and the supply vector ($\overrightarrow{U}$; black arrow) falls within the region bounded by the consumption vectors. **b** When the equilibrium is unstable (gray dot), two species cannot coexist even if the supply vector is within the bounded region. Which species wins depends on the initial density of the two species. We used the same parameters for these two plots.

Because two species that compete for essential resources are unlikely to meet the first and second rules, we can imagine that the bounded region for coexistence is narrow. We tested this by measuring the angle bounded by the consumption vectors when first and second rules are met. We found that both the experiment and the simulation showed that the bounded region was larger when two species competed for substitutable resources than for essential resources (Fig. S10).

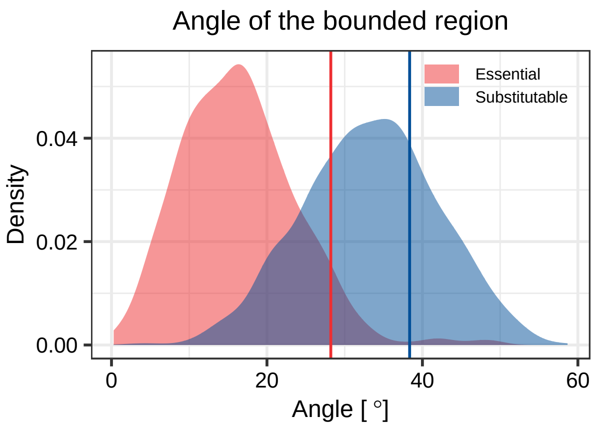


**Figure S10**: **The bounded region by consumption vectors is large when two species compete for substitutable resources.** Vertical lines indicate the angle of the bounded region calculated from the experiment. Density plots indicate the distribution of the angle calculated from the simulation (i.e., the theoretical expectations).

### Supplement S6 The case of multispecies community

Here, we scale up the Supplement S5 to multiple consumers.

#### Supplement S6.1 The first rule

The first rule should be adapted to: at least two of the species must be limited by different resources. In addition, the equilibrium of these two species must not be invadable by other species (i.e., all the other species have a negative growth rate at this equilibrium). Otherwise, at least one of the two species will be replaced by a third species. This condition is equivalent to: none of the species is "superior" (i.e., has the lowest $R^*$ for both resources). We can illustrate the case of competing for essential resources with Fig. S11a. The species 3 is "superior", having the lowest $R^*$ for resource 1 and 2. Although the ZNGIs of species 1 and 2 intersect, the equilibrium (intersect) can be invaded by species 3. Consequently, species 3 will outcompete species 1 and 2, irrespective of the resource consumption (the second rule). As long as none of the three species is a "superior" species, at least one uninvadable equilibrium is guaranteed (e.g., two in Fig. S11b and one in Fig. S11c). We can further see that this also applies to competition for substitutable resources in linear system (Fig. S11d-f). As we have shown in Supplement S5 that the nonlinear system was almost identical to the linear system regarding the first rule (as long as we ignore the case where two ZNGIs intersect twice, which is very rare). We can conclude that as long as none of the species has the lowest $R^*$ for both resources, the first rule is met, irrespective of the nonlinearity of the system.

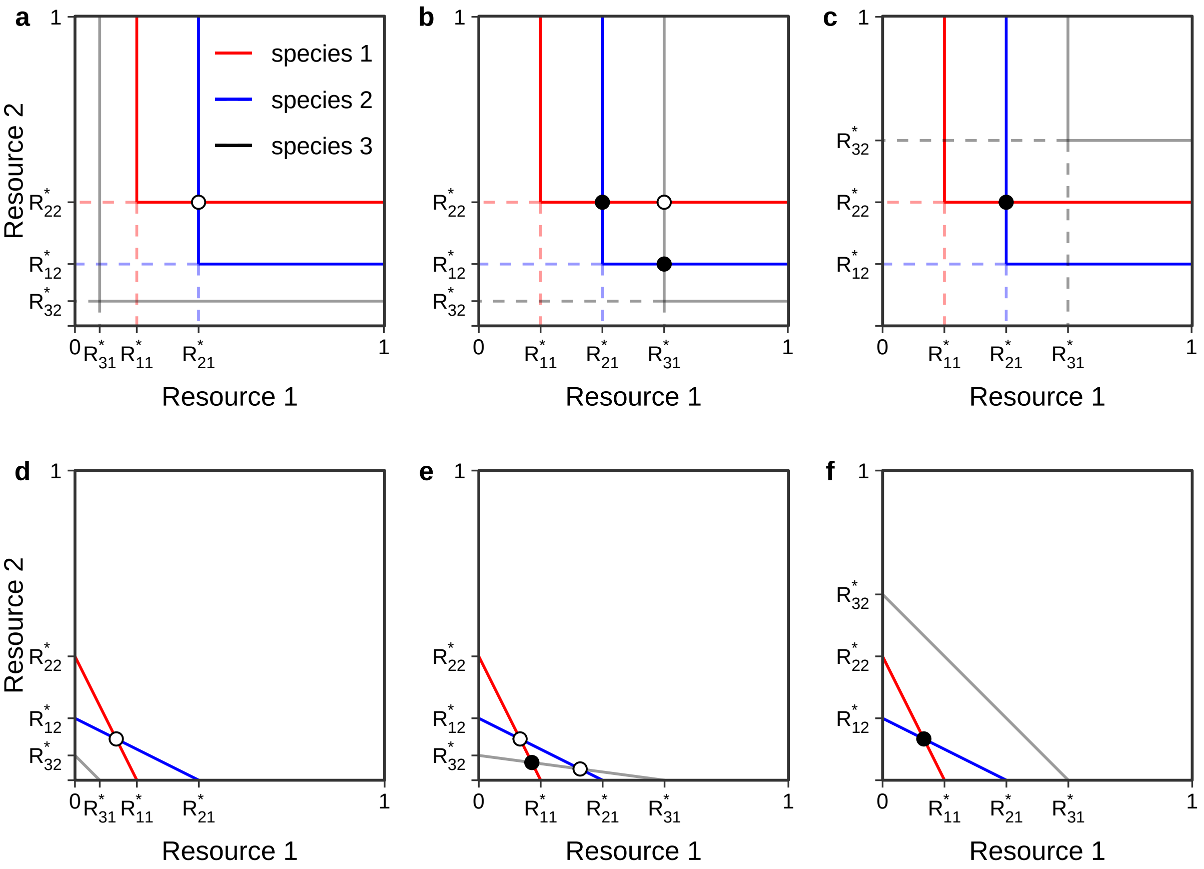


**Figure S11**: **Zero net growth isoclines (ZNGI) of three species competing for two resources (linear system).** **a, b, & c**, three species are competing for two essential resources. **d, e, & f**, three species are competing for two substitutable resources. We used the same parameters ($R^*$) of species 1 and 2 as Fig. SS6, and added a species 3 with different parameters. In **a & d**, species 3 is "superior", having the lowest $R^*$ in both resources 1 and 2. So, the first rule is not met. In **b, c, e, f**, none of the species is "superior". So, the first rule is met. We used the same parameters for the upper and the lower panels. Black dot indicates the equilibrium is uninvadeble and white dot indicates the equilibrium is invadable.

Since all $R^*_{ij}$ are independent (Supplement S5), the probability of none of the $S$ species is superior (i.e., has the lowest $R^*$ for both resources) is equivalent to the probability of rolling two fair $S$-sided dice and getting the different numbers on both. Here, each dice represents a resource and the number of the dice represents the species with the lowest $R^*$ on this resource. We can easily get the probability is $\frac{S-1}{S}$. This is further confirmed by the simulation of 999 communities (Fig. S12).

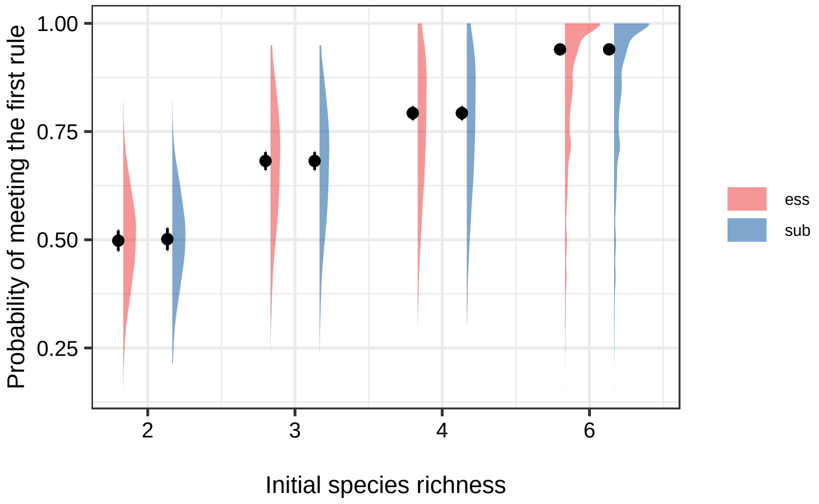


**Figure S12**: **Probability of the community meeting the first rule.** The data is from the simulation of 999 communities.

#### Supplement S6.2 The second rule

The second rule for multispecies communities is same as that for two-species communities. We only check it on the equilibrium that is uninvadable (i.e., black dots in Fig. S11). We were not able to solve it analytically. However, simulation showed that with increasing species richness, the probability of meeting the second rule increased (S13). This is because multiple non-invadable equiliria can be present in multispecies community (e.g., Fig. S11b and see Supplement S7 below).

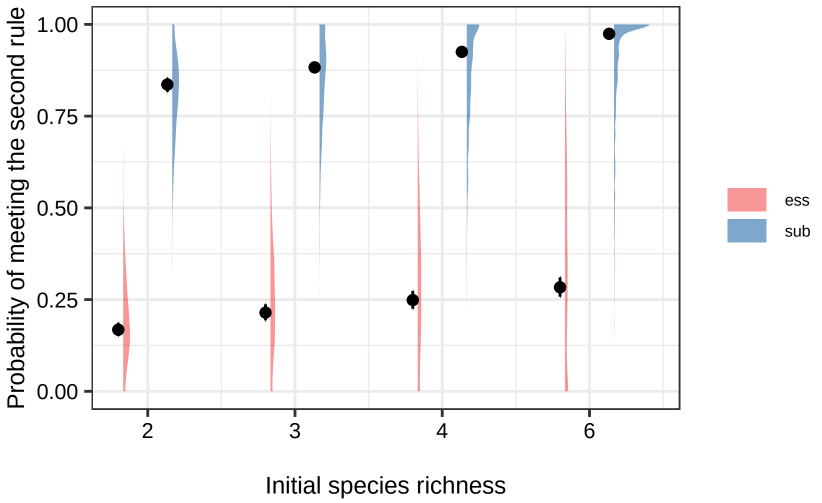


**Figure S13**: **Probability of the community meeting the second rule.** The data is from the simulation of 999 communities.

### Supplement S7 The presence of alternative stable states

When the first and second rules are met, a community can switch, with changing resource supplies, between coexistence, species 1 excluding species 2, and species 2 excluding species 1 (Fig. S9a). This is an example of alternative stable states. However, this type of alternative states is driven by the change of resource supply, which is not the case of our experiment (i.e., constant ratios of resource supply).

Still, there are two other types of alternative states, which do not rely on the change of resource supply. The first type includes two stable states of monocultures. It occurs when the first rule is meet but the second rule not. Under this condition, when the resource supply vector falls within the bounded region by the consumption vectors of two species, one will exclude the other, depending on the initial density (i.e., priority effect; Fig. S9b). As we have shown that the probability of community meeting the first rule rapidly increases with species richness, we can see that the probability of having alternative stable states increases with species richness [$P(meeting\ the\ first\ rule) \times P(not\ meeting\ the\ second\ rule)$].

The other type of alternative states includes multiple stable states of two species. It only occurs in multispecies communities when there are at least two uninvadable equilibria (e.g., Fig. S11b) and a certain resource supply vector must fall within at least two of the bounded regions at these equilibria. Again, we see that these type of alternative states increases with species richness (since it does not occur in two-species communities).

### Supplement S8 The model comparison (linear *vs.* nonlinear)

#### Supplement S8.1 The candidate models that were used to fit the data

We use three different models to fit the resource consumption and conversion with the monoculture experiment.

The first model assumed that both resource consumption and conversion were nonlinear (Equations 1 and 2 in the main text, hereafter the main model).

The second model assumed that resource consumption was nonlinear and resource conversion was linear (hereafter linear-conversion model):

$$\frac{1}{N}\frac{dR}{dt} = -\frac{cR}{s+R} = -X \tag{S22}$$

$$\begin{aligned}\frac{1}{N}\frac{dN}{dt} &= wX - m \\ &= \frac{cwR}{s+R} \\ &= \frac{u_{max}R}{k+R}\end{aligned} \tag{S23}$$

where $cw = u_{max}$ and $s = k$

The third model assumed that resource conversion was nonlinear and resource consumption was linear (hereafter linear-consumption model):

$$\frac{1}{N}\frac{dR}{dt} = -cR = -X \tag{S24}$$

$$\begin{aligned}\frac{1}{N}\frac{dN}{dt} &= \frac{wX}{q+X} - m \\ &= \frac{wcR}{q+cR} - m \\ &= \frac{wR}{\frac{q}{c}+R} \\ &= \frac{u_{max}R}{k+R}\end{aligned} \tag{S25}$$

where $w = u_{max}$ and $\frac{q}{c} = k$

#### Supplement S8.2 The performance

First, we extracted the R squared per species per resource for each model, we found that the main model and the linear-conversion model had higher median R squared (both 70.0%) than the linear-consumption model (63.9%).

Second, the same as we did for the main model, we predicted the community composition over time with the linear-conversion model and the linear-consumption model. We compared the predictive accuracy (on abundance and frequency) between the three models, with linear mixed effects models (as we did in the main text). We found that although all three models performed well, the main model outperformed the linear-consumption model (Fig. S14a & b; for abundance, $t$ = -4.41, $P < 0.001$; for frequency, $t$ = -2.42, $P$ = 0.02); and tended to outperform the linear-conversion model when predicting species frequency(Fig. S14b; $t$ = -1.47, $P$ = 0.143).

Together, these two results indicate that the main model (assuming nonlinear for both consumption and conversion) better quantified the resource requirement and consumption than the other two.

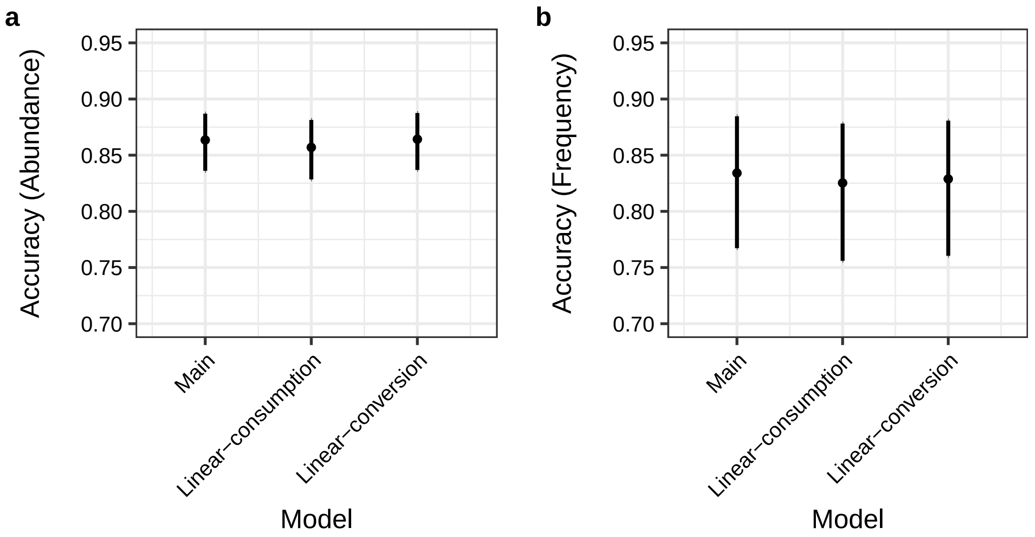


**Figure S14**: **Predictive accuracy of different models.** The main model assumed that both resource consumption and conversion were nonlinear. The linear-consumption model assumed that resource consumption was nonlinear and resource conversion was linear. The linear-conversion model assumed that resource consumption was linear and resource conversion was nonlinear.

### Supplement S9 Predicted resource concentration *vs.* measured resource concentration

Our study provided a novel method to predict resource concentration as long as the initial resource concentration was known. While it well-predicted community structure, it is unsure whether our predicted resource concentration matches the real resource concentration in the culture. To test this, we conducted an additional assay, where we grew *Chlamydomonas reinhardtii* for four days under three resource conditions (nitrate [N] = 512 uM, phosphorus [P] = 25.6 uM; N = 512 uM, P = 1.6 uM; and N = 32 uM, P = 25.6 uM). The setup is the same as the monoculture assay in the main text. For each resource condition, we set 16 replicates.

We first sampled 20 ul for each culture and measured the initial algal density with the IXM4. Then, from day one to day four, we collected a quarter of the replicates (i.e., 4 replicates) daily and filtered them through 0.2 um filters (Acrodisc@ Syringe filters, Pall). We pooled all the replicates from the same day for each resource condition to obtain a sufficient volume for measuring the resource concentration. Last, we quantified for each pooled sample the N and P concentrations with a spectrophotometer (Spectroquant® Prove 600, Merck), following the methods of Goldman & Jacobs [7] and Strickland [8]. In brief, to measure the N concentration, we added 1 ml diluted sample ($\times$ 1.5) into a 10 mm cuvette, added HCL 1N at 1:200 ratio, and measured the absorption at 220 and 275 nm. To measure the P concentration, we added 1 ml diluted sample into the 10 mm cuvette, added 0.1 ml reagent to avoid interferences with $CaCO_3$, and measured the absorption at 882 nm. To fit the calibration curve, we used fresh WC medium with different N concentrations (1, 2, 4, 8, 16, 32, 64, and 128 uM) and with different P concentrations (0.05, 0.1, 0.2, 0.4, 0.8, 1.6, 3.2 and 6.4 uM).

We found that in the fresh medium (used for the calibration curve), the detection limit for N was *c.* 8 uM, and for P, it was *c.* 0.4 uM. However, in the media where algae grew, the detection limit was much higher. For N, it was above 32 uM, and for P, it was above 1.6 uM. This is probably due to interference with small organic compounds in the media. Regardless of the exact reason behind it, the detection limit only allowed us to measure the N and P concentrations at day 1 and day 2 for the culture with the highest resource concentration (i.e., N = 512 uM and P = 25.6 uM).

We predicted the resource concentration (N and P) with the resource-consumer model, given the initial algal density. For comparison, we used models that assumed nonlinear consumption and conversion (main model), linear consumption and nonlinear conversion, and nonlinear consumption and linear conversion (section Supplement S8). We then compared the predicted resource concentrations with the measured concentrations.

We found that the measured resource concentration matched well with the resource concentration predicted by the main model (both consumption and conversion rate are nonlinear) or the linear-conversion model (Fig. S15). This indicates that our method well predicted the resource concentration and resource consumption.

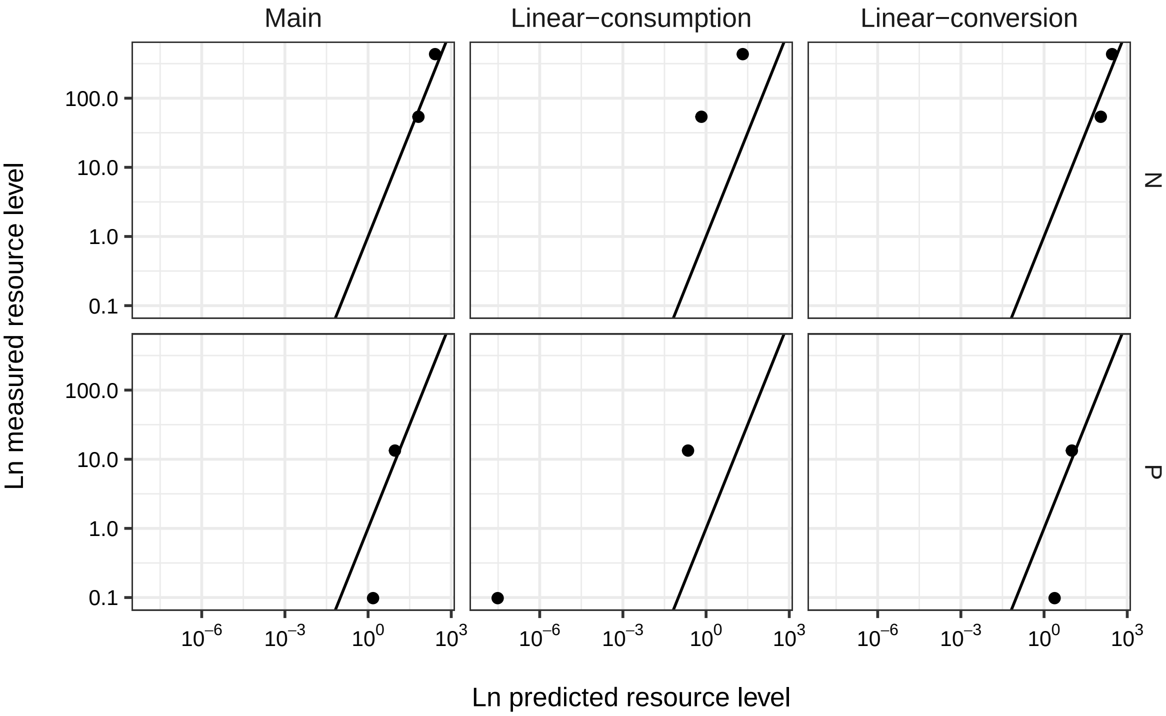


**Figure S15**: **Predictive accuracy of different models on resource level.** The main model assumed that both resource consumption and conversion were nonlinear. The linear-consumption model assumed that resource consumption was nonlinear and resource conversion was linear. The linear-conversion model assumed that resource consumption was linear and resource conversion was nonlinear. Y = X lines are plotted to enhance visualization. If the data points are on the lines, measured and predicted resource concentrations match. The unit of the concentrations is $\mu$M.

### Supplement S10 Imaging and machine learning

#### Supplement S10.1 Imaging and data acquisition

All samples were collected into 96-well plates, mixed with 120 ul WC medium, fixed with 2 ul of 1% PFA (paraformaldehyde) and 10% GA (glutaraldehyde), and settled overnight for imaging. We imaged each well at 15 sites (covering 68.66% area of the well) using three filter sets and the autofluorescence of the algal cells under a 10X magnification using IXM4 (ImageXpress® Micro 4 High-Content Imaging System). The three filters are Cy5 (exposure time: 10 ms), Cy3 (100 ms), and Texas red (100 ms), which excite the cell at three wavelengths and give us three images per site.

We used custom modules to detect cells in images (protocol will be provided upon acceptance). In brief, after Top-hat transform, the cell was detected based on the light intensity difference between the background and the cell in the Cy5 images. Then, we counted for each site the number of cells, and with the three wavelengths, exported 78 cell features that span autofluorescence intensity and cell morphology (e.g., cell length and size).

#### Supplement S10.2 Identifying species with machine learning

We built Neural Network Sequential Models to identify different algal species, using the *keras* [9] package in R. We randomly split the data (3.7 M cells) collected from the monoculture experiment and 196 additional monocultures (maintained over 12 days) into two parts: 80% of the cells were used as the training data and 20% as the test data. Since the predictive accuracy of models typically decreases with the number of classes (here, the number of species), we trained a separate model to classify each species combination in the competition experiment rather than training a single model to classify all 12 species.

For each model (i.e., species combination), we first filled the missing data with the mean of the corresponding feature. Second, we z-score transformed each of the cell features. Third, we built a Neural Network Sequential Model that contains four layers. The first to third layers contained 156, 60, and 30 units, respectively. The last layer outputted a length $n$ numeric vector (probabilities for each digit) using a softmax activation function, with $n$ equal to the number of species. Fourth, we assessed the accuracy of species identification with the test data. Last, we used the trained model to identify the species in the competition assay after transforming the cell features in the same way as the training data.

We found that although the classification accuracy declined with species richness (Fig. S16a; $F = 19.1, P < 0.001$), the overall accuracy of classifying different species was over 96% .

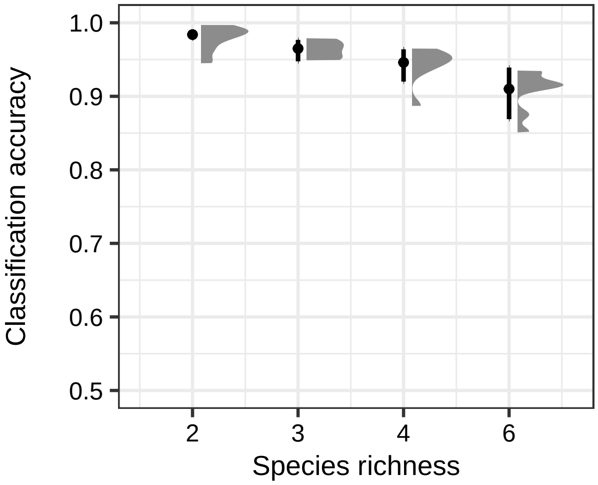


**Figure S16**: **Classification accuracy of the machine learning models.**

# Supplement S11    Growth rate as a function of two resources

To test how the algae grew under the limitation of two potential resources, we grew each of the 12 species for four days under eight different resource conditions. In four of them, we varied the concentrations of two essential resources, nitrate and phosphorus (red dots in Fig. S17a; $NO_3^-$ = 2 uM, P = 0.1 uM; $NO_3^-$ = 2 uM, P = 0.3 uM; $NO_3^-$ = 6 uM, P = 0.1 uM; $NO_3^-$ = 6 uM, P = 0.3 uM). In the other four, we varied the concentrations of two substitutable resources, nitrate and ammonium (red dots in Fig. S17b; $NO_3^-$ = 1 uM, $NH_4^+$ = 1 uM; $NO_3^-$ = 1 uM, $NH_4^+$ = 3 uM; $NO_3^-$ = 3 uM, $NH_4^+$ = 1 uM; $NO_3^-$ = 3 uM, $NH_4^+$ = 3 uM). The setup is the same as the monoculture assay in the main text. We set two replicates for each species per resource condition. This totaled 196 cultures. We sampled 20 ul for each culture on day one and day two and counted the algal abundance with the IXM4.

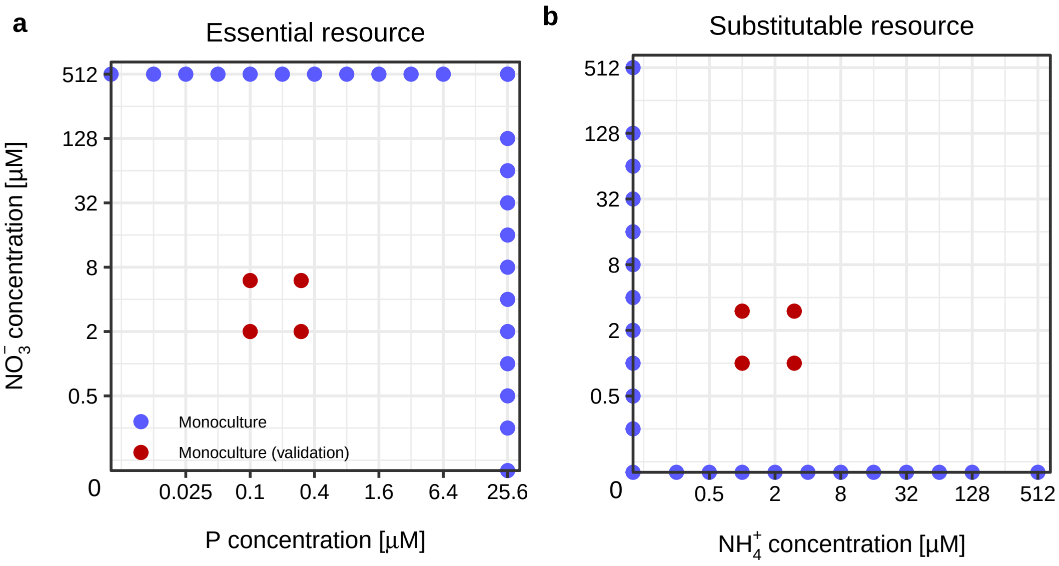


**Figure S17**: **The resource conditions of the monoculture experiments.** We conduct a monoculture experiment where the algae grew under the limitation of two essential resources (**a**) or two substitutable resources (**b**). Blue dots indicate the monoculture experiment where we quantified resource requirement and consumption. The red dots indicate the additional monoculture experiment where we assessed the growth rates.

We predicted the growth rate from day one to day two with two models. One assumed that the growth rate of algae was determined by the resource that supported lower growth rate (i.e., Liebig's law; equation S26). The other assumed that the growth rate was the sum of the growth rates provided by the two resources (hereafter, addition rule; equation S27):

$$\frac{1}{N_i}\frac{dN_i}{dt} = \min_{j=1,2}[\frac{u_{max,ij}R_j}{k_{ij}+R_j} - m_{ij}] \tag{S26}$$

$$\frac{1}{N_i}\frac{dN_i}{dt} = \sum_{j=1,2}[\frac{u_{max,ij}R_j}{k_{ij}+R_j} - m_{ij}] \tag{S27}$$

The function for resource consumption was kept the same as equation 4 in the main text, irrespective of the type of resources. After predicting the abundance on day two, we assessed the predictive accuracy with a Bray–Curtis similarity index, as we did in the main text:

$$acc = 1 - \frac{|a_{obs} - a_{pred}|}{a_{obs} + a_{pred}}$$

To assess whether the two models that assumed different resource types differed from each other, we conducted a linear-mixed effect model. The model included the predictive accuracy as the response variable; the type of resources (essential *vs.* substitutable) as the fixed effect; and the identity of medium and species as the random effects. The predictive accuracy was logit-transformed to improve the normality of the residuals. The Significance of the fixed effect was assessed with ANOVA. We did this for essential and substitutable resources separately.

We found that the model using Liebig's law outperformed the model using addition rule for species that competed for essential resources ($F_{1,176}$ = 6.96, P $<$ 0.001). In addition, the latter always over-predicted the algal abundance (the left panel in Fig. S18). For species that competed for subsitutable resources, it was the opposite ($F_{1,176}$ = 20.33, P $<$ 0.001). These support that the functions (equations S27 & S26) that we used for the growth rate were reliable.

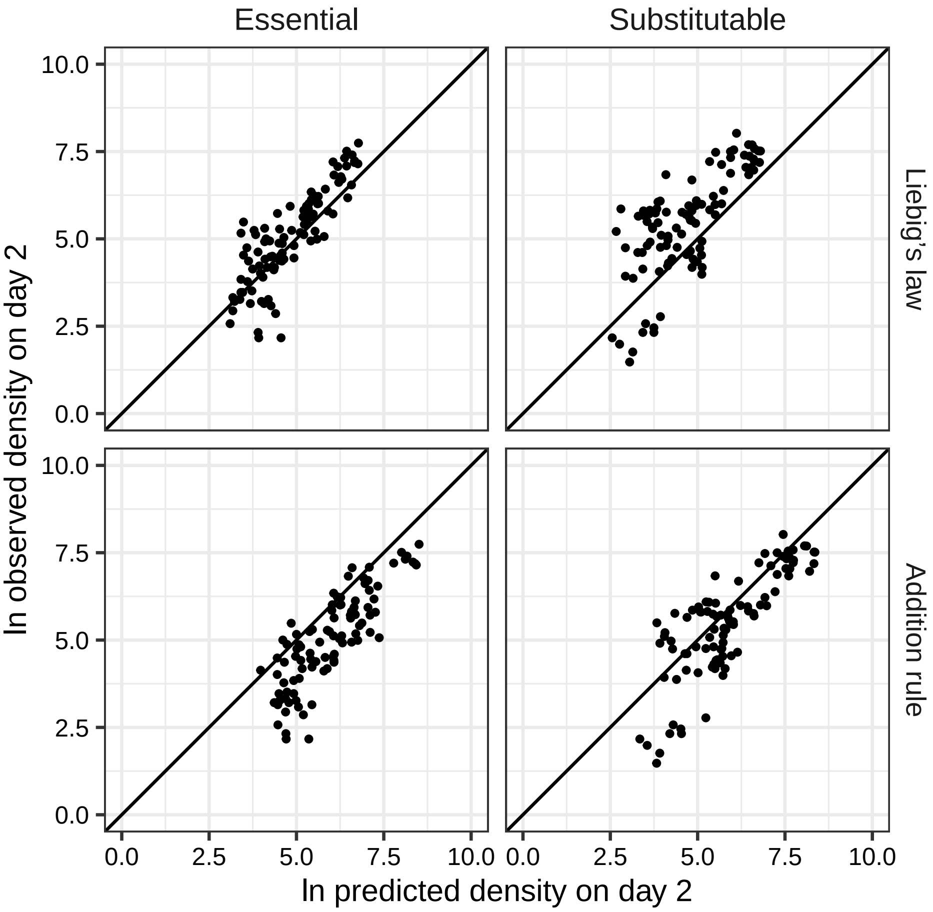


**Figure S18**: **The model accuracy.** Abundance on day two for each culture was predicted with models that either assumed Liebig's law or addition rule (the growth rate is the sum of growth rates on two resources). Y = X lines are plotted to enhance visualization. If the data points are on the lines, measured and predicted abundances match.
